# Reduction, removal or replacement of sodium nitrite in a model of cured and cooked meat: a joint evaluation of consequences on microbiological issues in food safety, colon ecosystem and colorectal carcinogenesis

**DOI:** 10.1101/2023.03.24.531666

**Authors:** Françoise Guéraud, Charline Buisson, Aurélie Promeyrat, Nathalie Naud, Edwin Fouché, Valérie Bézirard, Jacques Dupuy, Pascale Plaisancié, Cécile Héliès-Toussaint, Lidwine Trouilh, Jean-Luc Martin, Sabine Jeuge, Eléna Keuleyan, Noémie Petit, Laurent Aubry, Vassilia Théodorou, Bastien Frémaux, Maïwenn Olier, Giovanna Caderni, Tina Kostka, Gilles Nassy, Véronique Santé-Lhoutellier, Fabrice Pierre

## Abstract

**Scope:** Epidemiological and experimental evidence reported that processed meat consumption is associated with colorectal cancer (CRC) risk. Several studies suggest the involvement of nitrite or nitrate additives *via N*-nitroso-compound formation (NOCs).

**Methods and results:** Compared to the reference level (120 mg/kg of ham), the effects of sodium nitrite reduction (90 mg/kg of ham), removal and replacement were analysed on ham characteristics and in a CRC rat model. Sodium nitrite removal and reduction induced a similar decrease in CRC preneoplastic lesions, but only reduction led to (i) an inhibitory effect on *Listeria monocytogenes* growth comparable to that obtained using the reference nitrite level of 120 mg/kg and (ii) an effective control of lipid peroxidation. Among the three alternatives tested, none led to a significant gain when compared to the 120 mg/kg ham reference level: vegetable stock, due to nitrate presence, was very similar to this reference nitrite level, yeast extract induced a strong luminal peroxidation and no decrease in preneoplastic lesions despite the absence of NOCs, and polyphenol rich extract induced the clearest downward trend on preneoplastic lesions but the concomitant presence of nitrosyl iron in feces. Except vegetable stock, other alternatives were less efficient than sodium nitrite (≥ 90 mg/kg) in reducing *L. monocytogenes* growth.

**Conclusion:** Nitrite reduction (90mg/kg) effectively reduced CRC risk through limiting NOC formation and lipid peroxidation, while mitigating *L. monocytogenes* risks from cooked hams. Going further in reduction should be possible if accompanied by antioxidants to limit lipid peroxidation and appropriate use-by dates.

## 1. Introduction

In 2015, the state of epidemiological and experimental knowledge led the International Agency for Research on Cancer (IARC) to classify the consumption of processed meat (meat processed by salting, curing by nitrite or nitrate salts, fermentation, smoking, etc.) as carcinogenic to humans (Group 1, in particular for colorectal cancer risk (CRC)).^[1,2]^ The starting point for this classification was based on experimental and epidemiological findings obtained since the 1990s, which have highlighted a positive association between processed meat consumption and CRC risk (World Cancer Research Fund - WCRF, 1997).^[3]^ Since then, these first epidemiological data have been consolidated by meta-analyses, including those carried out by the World Cancer Research Fund.^[4]^ Moreover, experimental studies performed in a CRC animal model have demonstrated that consumption of cooked ham model and commercial hot dog sausages promoted colon preneoplastic lesions.^[5,6]^

In the context of human health, nitrite or nitrate salts and linked nitrogen species such as nitric oxide (NO) are the subject of growing scientific debate. Indeed, the ingestion of nitrate or nitrite salts under conditions that result in endogenous nitrosation is classified as "probably carcinogenic" to humans (Group 2A) by IARC from 2010.^[7]^ Beyond the mechanistic hypotheses involving heme iron, the carcinogenic properties of *N*-nitroso compounds (NOCs) are particularly suspected to explain the positive association between processed meat consumption and CRC risk.^[8]^ Nitrosation and nitrosylation, the two main reactions leading to the formation of NOCs in meat product and during digestion, are due to the frequent addition of the food additives E249 to E252 (potassium nitrite-E249, sodium nitrite-E250, sodium nitrate-E251 and potassium nitrate-E252) in processed meats.^[9]^ Nitrosation refers to the addition of a nitrosonium ion (NO^+^) to a nucleophilic group such as an amine, which generates nitrosamines or nitrosamides.^[10]^ Nitrosamines, which can be formed throughout the digestive tract either at acidic pH in the stomach or at neutral pH and in the presence of heme iron in the intestine, can lead to the formation of DNA adducts.^[11]^ Experimental toxicological data highlight the genotoxicity and carcinogenicity of these compounds. Among this important family of compounds, *N*-nitrosodimethylamine (NDMA) is identified as having the highest carcinogenic potential (SCCS 2011).^[12]^ The nitrosonium ion can also react with free thiol groups (R-SH), through an S-nitrosation reaction, to form nitrosothiols. In contrast, nitrosylation is a direct addition of NO to a reactant such as heme iron leading to the formation of nitrosyl heme.^[9]^

The nitrite and nitrate salts are used as food additives in various processed meat products in order to prevent or reduce the growth of pathogenic bacteria (e.g. *Clostridium botulinum, Clostridium perfringens, Listeria monocytogenes, Salmonella* spp.), to extend shelf life, to limit oxidation and to contribute to the color and taste of processed products (organoleptic functions).^[13]^ Regarding foodborne pathogens, Lebrun *et al*^[14]^ found that incorporation rates of sodium nitrite at a concentration of at least 30 mg/kg prevented the outgrowth and toxinogenesis of psychotropic *Clostridium botulinum* Group II type B in a cooked ham model. In comparison, the behavior of *L. monocytogenes* during the shelf-life of such product with reduced ingoing amounts of nitrite salt or alternatives to replace nitrite salt remains poorly described in the scientific literature; although a few studies have shown that added sodium nitrite may help control this pathogen in ready-to-eat meat products.^[15–17]^ *L. monocytogenes* is one of the most common foodborne pathogens and its detection in the processed meat products accounted for 32% of the recalls registered for these products in the last year in France (data collected on the *rappel.conso.gouv* website of the French government). Concerning carcinogenesis, several experimental and epidemiological studies conducted specifically on the food additives nitrate and nitrite suggested a role of food additive-induced nitrosation and nitrosylation in CRC promotion. Indeed, regarding epidemiology, two meta-analyses had recently highlighted positive associations between nitrate salts (but not nitrite salt) from the diet as a whole and the risk of colorectal^[18]^ and ovarian cancer.^[19]^ But authors did not distinguish between natural nitrite/nitrate salts from water, vegetables and nitrite/nitrate salts supplied as food additives. However, a recent study conducted on the NutriNet-Santé cohort allowed to distinguish the different dietary sources of nitrate and nitrite salts (natural food sources or food additives) and their respective association with cancer risk.^[20]^ Compared to non-consumers, high consumers of nitrated and nitrited food additives had a higher risk of several cancers: the food additives nitrate and nitrite salts were positively associated with breast and prostate cancer risks, respectively, while no association was observed for nitrite/nitrate from natural sources. Regarding experimental data, we previously demonstrated that a diet based on cooked meat product without nitrite salt limited the promotion of CRC in a rodent model pre-treated with azoxymethane as opposed to a diet composed of cooked processed meat with 120 mg/kg of sodium nitrite.^[21]^ Despite this clear effect on CRC promotion in this animal model, the absence of sodium nitrite was associated in this study to an increased endogenous lipid peroxidation along with a high toxic alkenal (such as 4-hydroxynonenal (HNE)) absorption as reflected by increased urinary di-hydroxynonane mercapturic acid (DHN-MA) excretion.^[21]^ The production and absorption of those toxic alkenals could induce deleterious effects in extra-intestinal locations as proposed by Boléa *et al*.^[22]^

Interestingly, these studies in a rat model did not demonstrate any association between CRC promotion, consumption of processed meats and total NOCs ^[23,24]^, but highlighted a positive association with a subcategory of NOCs: the nitrosylated heme iron originated from the chemical reaction between heme iron and nitrite salt. Nitrosyl heme is formed within the processed meat products (responsible for the typical pink color of cooked processed meats) but also during digestion in the small intestine. Indeed, due to the alkaline conditions in the lower part of the gastrointestinal tract, *S*-nitrosothiols are degraded and the released nitric oxide may also react with heme to form nitrosyl heme.^[25]^ The numerous studies on nitrosamines carcinogenicity and our previous studies proposing an association, in the rat model of CRC, between nitrosylated iron and CRC development highlight the need to limit population exposure to these food additives.

This study provides an original multidisciplinary approach considering food safety, technological properties and toxicological aspects to examine the effect of reducing or removing ingoing sodium nitrite, but also of current or future nitrite salt replacement strategies including antioxidants, vitamins and natural phenolic compounds on (i) endogenous reactions (nitrosation, nitrosylation and peroxidation), (ii) colonic ecosystem (microbiota and colon mucosal detoxification capacities), (iii) promotion of colon carcinogenesis in a CRC rat model, (iv) growth potential of *L. monocytogenes* on a sliced cooked ham model. This study should provide relevant and useful highlights to food regulation agencies and to implement appropriate short-term strategies for nitrite reduction or replacement in the meat processing sector.

## 2. Experimental section

### 2.1. Production, microbiological and biochemical characteristics of cooked ham model products

#### 2.1.1. Production of cooked ham model products

The experimental cured meat, similar to air-exposed model ham and called DCNO (for Dark Cooked meat with sodium Nitrite, Oxidized) in our previous studies, was chosen because it promotes colon carcinogenesis in rats.^[21]^ It was used as the reference and positive control in this study to test potentially protective effects of reduction, removal or alternatives to sodium nitrite. It was produced with the usual sodium nitrite concentration in France, *i.e.* 120 mg/kg. Dark red meat was obtained from fresh defatted, derinded, denerved and deboned pork shoulders. The product without sodium nitrite was the negative control of this study.

For the carcinogenesis study, six experimental cooked ham models were made in a ham factory according to a process allowing a homogeneous curing treatment. Each model is composed of 80 kg ground pork (10-12 mm) mixed with appropriate concentrations of ingredients/additives as describe in Table S3 of Supplementary Material. Three concentrations in sodium nitrite were added in brine, at the final rate in finished products: 120 mg/kg (Ni- 120) the maximum authorized by the French Code of Practice, 90 mg/kg (Ni-90) for the reduced level and 0 mg/kg (Ni-0) for the cooked ham model without sodium nitrite. Three alternatives were added in the brine without sodium nitrite, at a concentration recommended by suppliers: a vegetable stock (VS) that contained different ingredients: sugar, a powdered juice of chards, dehydrated carrot (Bouillon Nat 223, DAT-Schaub France, Thiais, France), an aroma rich in polyphenol and ascorbic acid called “polyphenol-rich extract” (PRE), and a yeast extract (savor lyfe NR01) and *Staphylococcus xylosus* culture (Lalcult Carne Rose 01) from Lallemand SAS, Blagnac, France) called YE. After curing, the ground pork was tumbled for 18h at 2 700 rotations. Then, the cured ground pork was transferred into moulds and packed into heat shrink bags under vacuum condition. Vacuum packed products were steam cooked up to a core temperature of 68°C for 990 min, with a temperature stage at 50°C for 60 min, for the development of both the YE and VS starters.

After this thermal processing, 685 min were required to fall the temperature from 67°C to 4°C (representative of a pasteurization value P70/10 of 90 min). Cooked ham models were unpacked, sliced into a 15 mm thickness and 60 kg of sliced cooked ham were packed in 350 g top sealed trays under protective atmosphere (50 % CO_2_ and 50 % N_2_). Some of these samples were stored at −20°C (D02: day 2 post packaging) for the carcinogenesis study and the other ones were stored 14 days at 4°C and 35 days at 8°C for the biochemical analyses.

#### 2.1.2. Biochemical characteristics of cooked ham models

The samples were frozen before being ground in liquid nitrogen to avoid any oxidation before analysis.

##### 2.1.2.1 Iron, heme iron, nitrosylheme and lipid peroxidation

Total heme iron, lipid peroxidation, nitrosylheme content and concentration of total non-heme iron, Fe^2+^, and Fe^3+^ were assessed as previously described.^[9]^ More details are supplied in Supplementary Materials.

##### 2.1.2.2. Residual nitrite, residual nitrate, nitrosothiols and non-volatile nitrosamines

Nitroso-compounds (total non-volatile *N*-nitrosamines and nitrosothiols), residual nitrite and nitrate were quantified as described previously.^[9]^ More details are supplied in Supplementary Materials.

### 2.2. Growth potential of *Listeria monocytogenes* in cooked ham model products

Growth potential of *L. monocytogenes* was followed at different sampling times during shelf-life of sliced cooked ham model products, including 6 different recipes (see Supplementary Material and Table S3). Three specific and independent batches of sliced cooked ham model products were used for this assay. They were produced following similar processes to those employed in the meat processing industry. A cocktail of three *L. monocytogenes* strains in equal proportions was prepared in saline solution, which was used to surface inoculate slices of the different cooked ham model products. These samples were then stored for 14 days at 4 °C followed by 35 days at 8 °C. Especially, three sliced cooked ham model samples per recipe and batch were analyzed for the enumeration of *L. monocytogenes* using the BRD 07/17 - 01/09 standard method at each sampling date (D0, D07, D14, D17, D22, D28, D35 and D49) (see Supplementary Material for more details).

### 2.3. Study of carcinogenesis: animals, design and diets

#### 2.3.1. Ethical approval

Animal experiment was approved by the local Ethical Committee (CE n°86), authorized by the French Ministry of Research (APAFIS 27180_2020091017209910_v2) and conducted in accordance with the European Council on Animals used in Experimental Studies and ARRIVE guidelines.

#### 2.3.2. Animals, design and diets

Male Fischer 344 (F344/DuCrl) rats were purchased from Charles River Laboratories, 12 rats/group, aged 5-6 weeks. After acclimatization, they received a single i.p. injection of azoxymethane (Sigma; 20 mg/kg) in NaCl (9 g/L water) to induce colon preneoplastic lesions. Seven days later, the rats were randomly assigned to six groups and fed the experimental diets daily *ad libitum* (at the end of the afternoon in order to avoid any important oxidative degradation before food intake) for 98-99 days before euthanasia. Colons were then removed, washed, opened, coded, and fixed in 10% buffered formalin (Sigma-Aldrich) before Mucin Depleted Foci (MDF) analysis. Body weight was monitored every week during the 4 first weeks, then every two weeks. Food and water intakes were measured on day 15 and day 85. Feces were collected during 3 days on days 72 to 74 or 79 to 81. Each rat was placed in a metabolic cage and urine was collected on day 72 or on day 79. Anal feces for microbiota analysis were collected at the end of the experimental diet period and kept at −80°C before analysis. Fecal and urine samples were kept at −80°C and −20°C, respectively before analysis.

Experimental diets were based on powdered low-calcium, no-fat AIN-76 rodent diets (SAAJ, Jouy-en-Josas, France), supplemented with 5% safflower oil (MPBio), with 45% (dry weight) of experimental cured meats and 8% of gelatin. Model cooked ham trays were opened and stored at 4°C for 3.5 days before rat diet preparation (every two weeks), to induce an oxidation step related to bad storage conditions, the diets were then prepared by mixing ham, the AIN-76-based powder and the oil, and then frozen at −20°C until use.

#### 2.3.3. Preneoplastic lesions scoring

Formaline-fixed colons were stained with high iron diamine-Alcian blue procedure (HID-AB)^[26]^ for scoring the preneoplastic lesions MDF. Scoring was achieved by a reader blinded for colon sample origin.

### 2.4. Urinary, fecal, and microbiota analysis

#### 2.4.1 Urinary 1,4-dihydroxynonane mercapturic acid (DHN-MA)

DHN-MA, the major urinary metabolite of HNE^[27]^ was measured using a Bertin Bioreagent kit (Montigny-le-Bretonneux, France). Urines were diluted (1/1000) in the kit buffer.

#### 2.4.2 Fecal water preparation and analysis of heme, TBARS and fecal activities

Fecal waters were prepared from feces collected during 24h. Feces were added with ultrapure water (0.5 g/mL). To prevent oxidation, 50 µl/mL butylated hydroxytoluene (BHT) was added to water. The feces mixture was homogenized in Lysing Matrix E tubes and FastPrep technology (MPBiomedicals) 3 times for 30 seconds (6 m/s) and then centrifuged at 3 200 g for 20 minutes at 4°C, as described previously, with slight modifications.^[28]^ Supernatants (fecal waters) were collected and kept at −80°C until use. Heme was measured with the Heme Assay (Kit MAK316 – Sigma) according to the recommendations of the manufacturer; fecal TBARS were measured in fecal waters according to Ohkawa *et al*.^[29]^ For cytotoxicity and genotoxicity, fecal waters were tested on normal immortalized colon epithelial cells (Co cells), which were established as described previously.^[30,31]^ Conditionally immortalized cells were seeded into 96-well plates at 15×10^3^ cells per well in permissive conditions. After reaching sub confluence (70%), cells were transferred to 37°C and treated for 24h with filtered fecal waters in serum-free DMEM to avoid any reaction between serum and fecal water. Cytotoxicity was assessed using WST-1 kit as described earlier ^[28]^ and genotoxicity was measured using the ɣ H2AX in-cell Western blot assay according to Khoury *et al*.^[32]^

#### 2.4.3 Quantification of fecal *N*-nitroso compounds (NOCs)

Fecal water aliquots were transferred in amber reaction tubes and stored at −80°C until further use. The following steps and analyses were done under reduced light conditions, while all samples and chemicals were kept on ice. Fecal water samples were analysed for NOCs using an Ecomedics CLD88 NO-Analyzer (Eco Physics GmbH, Hürth, Germany) as published recently with slight modifications.^[33]^ See supplementary material for more details.

#### 2.4.4 High-throughput 16S rRNA gene amplicon analysis

Genomic DNA from snap-frozen fecal samples and the amplification of hypervariable V3-V4 regions of their 16S rRNA gene were prepared as described previously.^[34]^ Libraries preparation and sequencing (Illumina Miseq cartridge) were performed by the Genotoul facility (Get-Biopuces, Toulouse). Raw sequences were processed using the FROGS pipeline (Galaxy Version 3.2.3) and analyzed using the R package Phyloseq (v1.34.0) as follows ^[35]^: Each pair-end valid denoised sequences were filtered, merged and clustered with the swarm fastidious option using a maximum aggregation distance of 1.^[36]^ Putative chimera were removed (vsearch) and clusters (i) whose abundance represented at least 0,005 % of all sequences, (ii) presents in at least 2 times in a minimum of 5% of total samples with a prevalence threshold of 5% of all samples, were retained, yielding to 318 final clusters. The silva 138.1_16S reference database was used for cluster affiliation into Operational Taxonomic Units (OTUs) using Blast+with equal multi-hits, taxonomic multi-affiliations were checked manually. Within sample community richness and eveness (α diversity) were estimated using both the Chao-1 and Shannon indexes respectively, and examined by one-way ANOVA analysis and Tukey’s multiple comparisons test. Divergence of bacterial composition between samples (β diversity, Phyloseq) was explored using the Unifrac distance matrices, and statistically tested using permutational multivariate analysis of variance (MANOVA using Adonis test with 9999 permutations followed by pairwise multilevel comparisons). OTUs were agglomerated at the order and genus ranks to further analyze differential abundances according to diets using the Deseq2 package (v1.30.1). Detailed results of statistical analyses were reported in Table S10 as supporting information. For graphical visualization purpose in parallel, raw 16S counts of taxa, whose abundance was significantly affected by diets, were normalized according to the mixMC pipeline.^[37]^

### 2.5 Statistical analyses

For the animal experiment, once outliers were identified/removed by a ROUT test, and normality was checked, ANOVA was performed using GraphPad Prism software (version 9.5.0), followed by a Dunnett’s or a Tukey’s mean comparison test, in case of comparison to a control group (sodium nitrite effect), or comparison of all means (alternative effect), respectively. In case of heteroscedasticity (tested by Bartlett’s and Brown-Forsythe tests), data were log transformed before ANOVA (DHN-MA and NOCs). In case of non-normality of residuals, a nonparametric test (Kruskal-Wallis) was performed followed by Dunn’s mean comparison test (some RT-qPCR and bacterial taxa). Finally, if equal SD cannot be assumed despite data transformation, a Welch’s ANOVA was performed (cytotoxicity test). The growth potential values of *L. monocytogenes* were compared among the different recipes using the ANOVA test followed, when significant, by pairwise comparison using the estimated marginal means (Emmeans test) followed by Bonferroni correction. Two-side analyses were used throughout, and p values less than or equal to 0.05 were considered significant.

## 3. Results

### 3.1. Impacts of sodium nitrite concentrations on cooked ham model products, microbiological risk, colon ecosystem and colorectal carcinogenesis

#### 3.1.1. Consequences of sodium nitrite concentrations on nitrosylation, nitrosation and lipid peroxidation in the cooked ham model

Compared to the reference cooked ham model produced with 120 mg/kg of sodium nitrite (Ni-120), the reduction of the food additive to 90 mg/kg (Ni-90) did not induce a decrease of nitrosylation of heme iron, with an equal percentage of nitrosylation. In the same way, reduction from 120 to 90 mg/kg did not result in a significant modification of total non-volatile *N*-nitrosamines and nitrosothiols but induced however a significant reduction (p < 0.05) of residual nitrite and nitrate (Table 1). Compared to Ni-120, the removal of the food additive resulted in a strong and significant decrease of nitrosylated iron in the cooked ham model with a percentage of nitrosylation falling from 58 to 9%. The absence of added sodium nitrite also induced a disappearance of nitrosothiols, residual nitrite and a significant decrease in residual nitrate but a strong increase of lipid peroxidation (TBARS) in comparison to Ni-120 (Table 1). Change from 90 to 0 mg/kg of sodium nitrites reduced also significantly the nitrosylation of heme iron, the presence of non-volatile *N*-nitrosamines, residual nitrite and nitrate and increased lipid peroxidation (p < 0.05; Table1).

**Table 1:** Impact of sodium nitrite levels on biochemical characteristics of cooked ham models (0 *vs* 90 *vs* 120 mg/kg). Mean values with unlike letters were significantly different (p ≤ 0.05). Data are mean ± SD.

|  | Ni-120 | Ni-90 | Ni-0 |
| --- | --- | --- | --- |
| **Total heme iron ($\mu\text{M}$)** | 189 ± 40 <sup>$\alpha$</sup> | 146 ± 5 <sup>$\alpha$</sup> | 111 ± 9 <sup>$b$</sup> |
| **Nitrosylated iron ($\mu\text{M}$)** | 110 ± 11 <sup>$\alpha$</sup> | 83 ± 6 <sup>$b$</sup> | 10 ± 1 <sup>$c$</sup> |
| **% Nitrosylated iron** | 58 | 57 | 9 |
| **TBARS ($\mu\text{g MDA/g}$)** | 0.041 ± 0.006 <sup>$\alpha$</sup> | 0.042 ± 0.004 <sup>$\alpha$</sup> | 0.303 ± 0.002 <sup>$b$</sup> |
| **Total free iron (mg/kg)** | 4.92 ± 0.04 <sup>$\alpha$</sup> | 6.45 ± 0.04 <sup>$b$</sup> | 5.97 ± 0.01 <sup>$c$</sup> |
| **Total non volatile *N*-nitrosamines (mg/kg)** | 2.09 ± 1.02 <sup>$ab$</sup> | 4.08 ± 0.28 <sup>$\alpha$</sup> | 1.27 ± 0.17 <sup>$b$</sup> |
| **Nitrosothiols (mg/kg)** | 2.05 ± 0.25 <sup>$\alpha$</sup> | 1.38 ± 0.21 <sup>$\alpha$</sup> | 0.00 ± 0.00 <sup>$\alpha$</sup> |
| **Residual nitrites (mg/kg)** | 9.36 ± 0.23 <sup>$\alpha$</sup> | 6.76 ± 0.11 <sup>$b$</sup> | 0.00 ± 0.00 <sup>$c$</sup> |
| **Residual nitrates (mg/kg)** | 24.95 ± 1.07 <sup>$\alpha$</sup> | 14.59 ± 0.97 <sup>$b$</sup> | 4.51 ± 0.01 <sup>$c$</sup> |

#### 3.1.2. *Listeria monocytogenes* growth in sliced cooked ham model products as function of sodium nitrite levels

Under the storage conditions tested, *L. monocytogenes* was able to grow in sliced cooked ham model samples with added sodium nitrite salt up to 120 mg/kg (δ > 0.5 Log_10_ CFU/g by day 14 for the three tested recipes, Figure 1). However, growth of the pathogen was significantly reduced during the first 3 weeks in samples prepared with 90 or 120 mg sodium nitrite per kg of meat compared to those without added nitrite salt (p ≤ 0.01). Growth potentials were not significantly different between samples containing 90 or 120 mg NaNO_2_/kg within this period (p > 0.05). After 28 days of storage, growth potentials were similar between all recipes, regardless of the ingoing amounts of nitrite salt employed.

**Figure 1.**
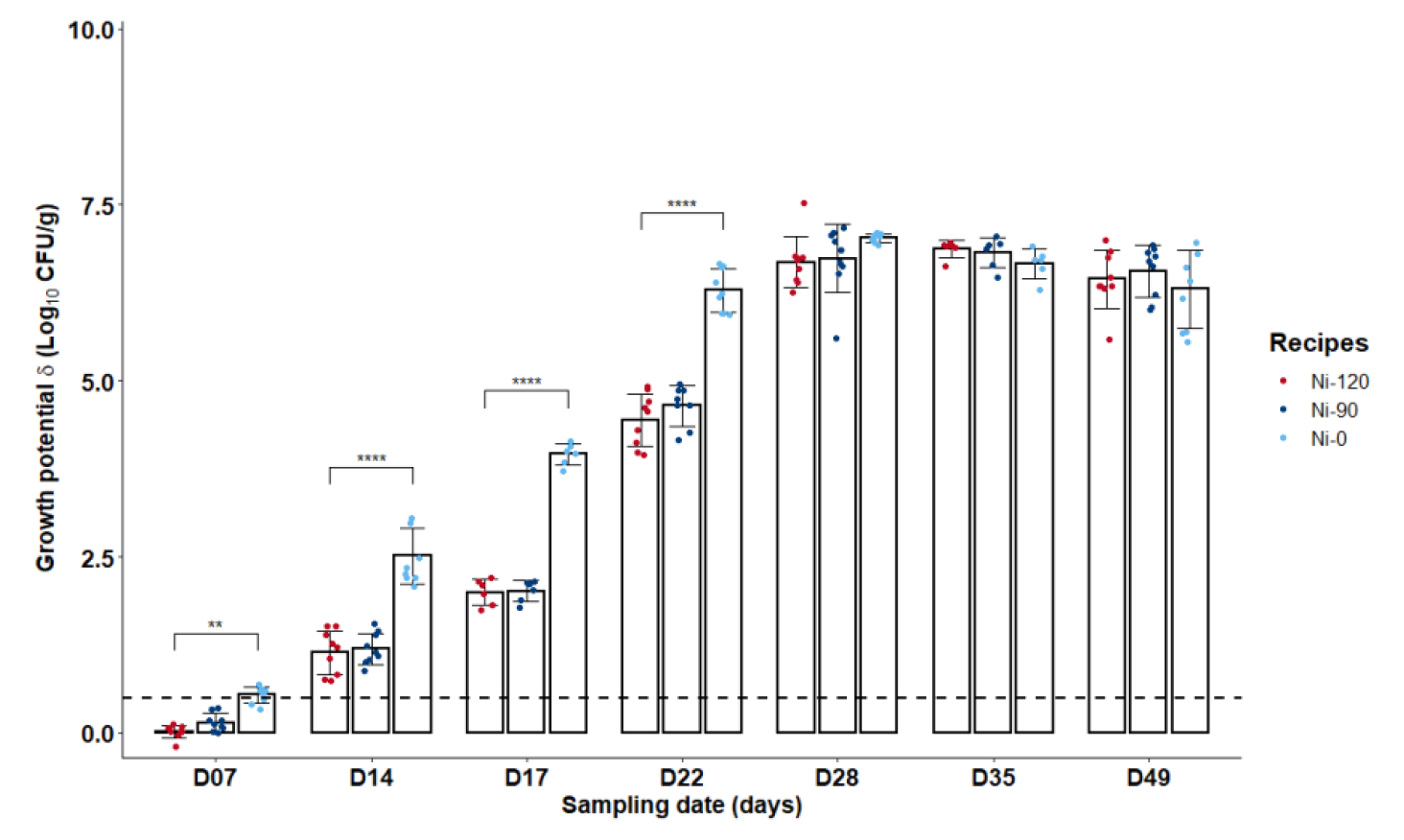
Growth potentials (δ) of *Listeria monocytogenes* at several sampling times during shelf-life of the sliced cooked ham model products with different sodium nitrite levels (0 *vs* 90 or 120 mg/kg). Data were represented using scatter plots with bar (mean ± SD), **p ≤ 0.01, ****p ≤ 0.0001. The dashed line represents the limit value of 0.5 Log_10_ CFU/g above which the cooked ham model samples support the growth of *L. monocytogenes*.

#### 3.1.3. Sodium nitrite removal or reduction had strong impacts on endogenous formation of nitroso compounds, luminal and global peroxidation but not on cytotoxic and genotoxic activities of fecal waters

The reduction from 120 to 90 mg/kg and removal of sodium nitrite induced a dose-dependent decrease in the formation of total nitroso compounds (ATNC) and the three subcategories measured in the rat feces. Compared to the reference dose (Ni-120, the reduction (Ni-90) indeed induced a significant decrease (p ≤ 0.05) for 3 biomarkers of endogenous nitrosation and nitrosylation (ATNC, FeNO and RNNO) and the removal induced a near disappearance of these fecal markers with an almost total absence of fecal total NOCs, nitrosylated iron, RNNO and RSNO (Figure 2A). Conversely, we observed a dose-dependent significant increase in luminal lipid peroxidation measured in fecal waters (Figure 2B, TBARS) and a dose-dependent increase in urinary excretion of DHN-MA (Figure 2B), the major metabolite of HNE indicating an increase in HNE absorption, probably due to an abundant presence of this lipid oxidation product in intestinal lumen. These consequences of sodium nitrite reduction or removal are not associated with higher levels of heme iron, a peroxidation catalyst (Figure 2C). These modifications in the fecal contents of *N*-nitroso compounds and alkenals were not associated with a change in the cytotoxic and genotoxic activities of fecal waters (Figure 2D & E).

**Figure 2.**
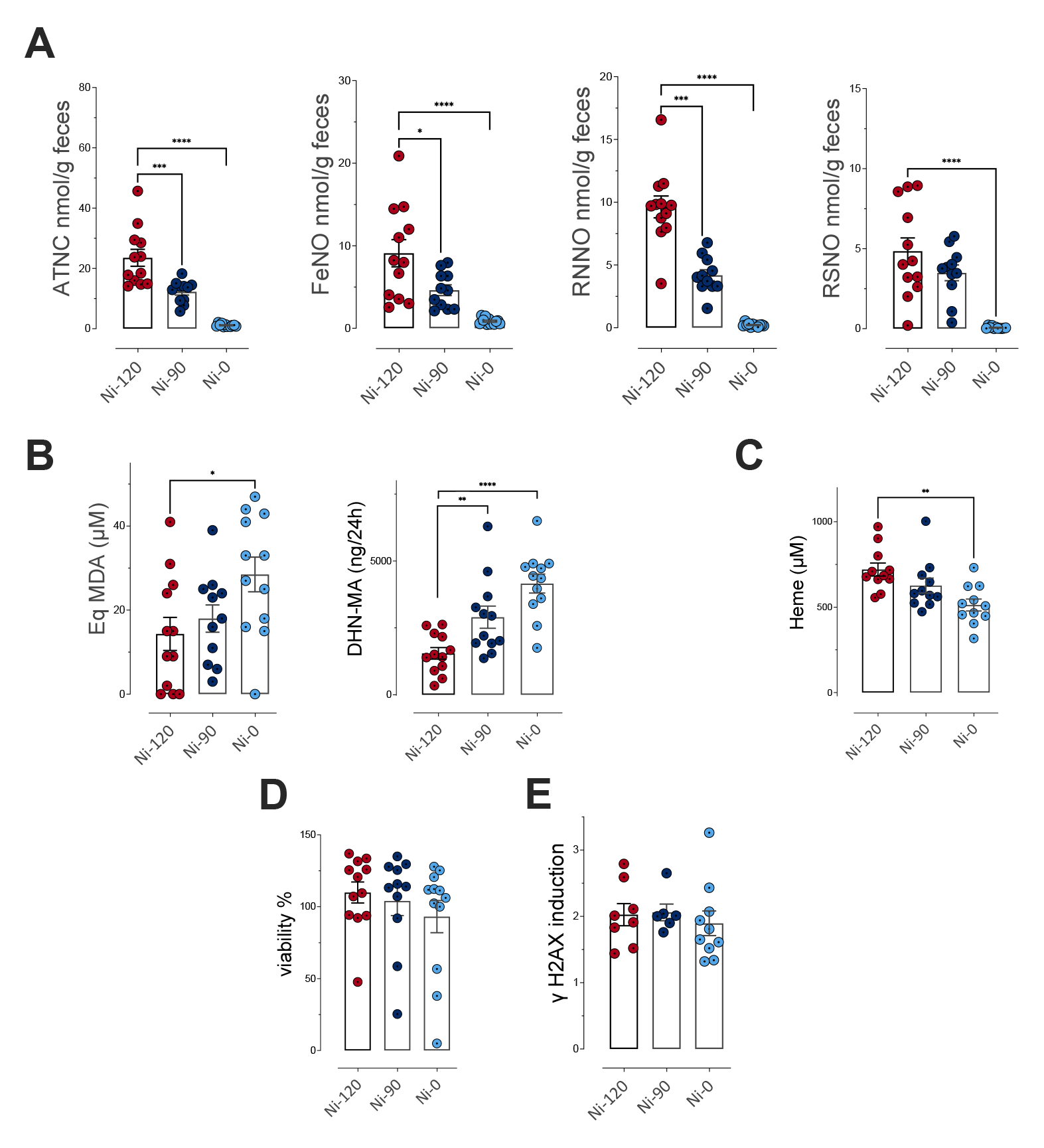
Impact of sodium nitrite levels in cooked ham models (0 *vs* 90 *vs* 120 mg/kg) on fecal and urinary biomarkers of lipid peroxidation and NOC formation. **A**-Nitroso-compounds in fecal water measured as apparent total NOCs (ATNC), as nitrosyl iron (FeNO), *N*-nitroso compounds (RNNO) and S-nitrosothiols (RSNO) (nmol/g of feces). **B**-Lipid peroxidation measured as TBARS (MDA equivalents, µM) in fecal water and DHN-MA in urine of 24h (ng/vol of 24h). **C**-Heme in fecal water (µM). **D**- Cytotoxic activity of fecal water as % of cellular viability. **E**- Genotoxic activity of fecal water. Data were represented using scatter plots with bar (mean ± sem), * p ≤ 0.05; **p ≤ 0.01; ***p ≤ 0.001.

#### 3.1.4. Effect of sodium nitrite levels in processed meats on the community distribution and diversity of the fecal microbiota as determined by 16S rRNA gene sequencing

Compositional analysis performed at the order level (Figure 3A), as well as diversity characterization at a finer level (Figure 3B and 3C), did not reveal major changes in the fecal microbiota in response to sodium nitrite reduction (Ni-90) in cooked ham model diet. By performing differential analysis at the order level, however, significant alterations in *Peptococcales* and *Peptostreptococcales-Tissierellales* were detected in the microbiota of rats fed the processed meat in which sodium nitrite was removed (Ni-0) (Figure 3A). At the genus level (Figure 3D), 6 bacterial communities underwent significative variations with sodium nitrite content and showed that reduced sodium nitrite intake was associated with a decrease in *Candidatus Soleaferrea*, *Romboutsia*, and genera belonging to the *Ruminoccocus torques* group, and an enrichment of *Eisenbergiella* and unknown genera belonging to *Peptococcaceae* and *Lachnospiraceae* (Figure 3E).

**Figure 3.**
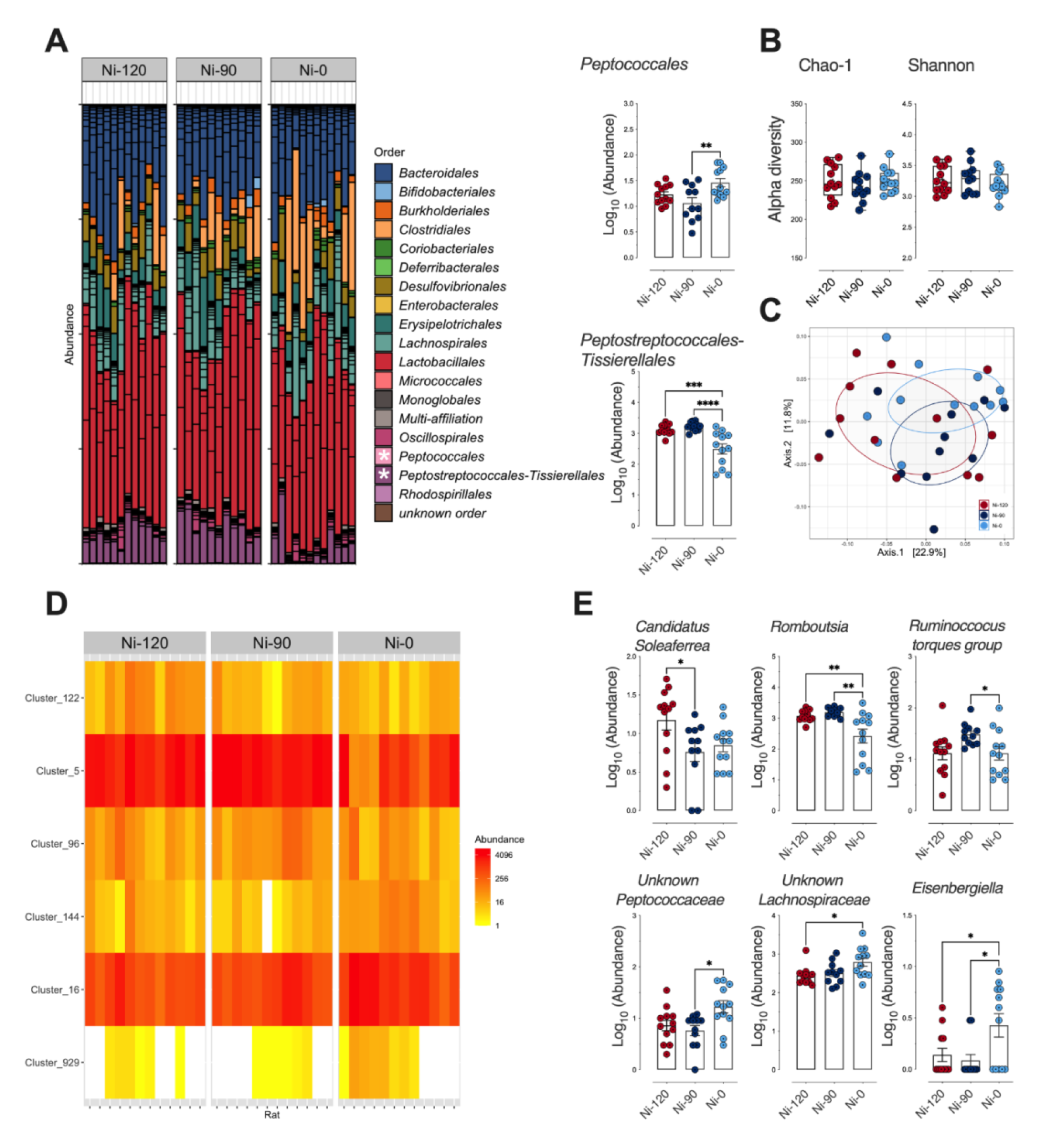
Impact of sodium nitrite levels in cooked ham models (0 *vs* 90 *vs* 120 mg/kg) on fecal microbiota. **A**-Distribution of bacterial communities at the order level. * Significant impact on *Peptococccales* and *Peptostreptococcales-Tissierellales* using differential abundance analysis at the order level (Deseq2, *P*_adj_≤0.05): Normalized Log_10_ abundances were represented using scatter plots with bar (mean ± sem). **B**-Alpha diversity: No significant impact seen on richness (Chao-1) or eveness (Shannon). Individual values are represented using box and whiskers (+ mean). **C**-Beta diversity (Unifrac distances, manova p>0.05). No significant difference between microbiota of rats fed the 3 cooked ham model diets. **D**-Heatmap of clusters agglomerated at the genus level affected by sodium nitrite content in ham-based diets. *P*_adj_≤ 0.05 using differential abundance analysis (Deseq2). **E**-Normalized abundance of the 6 clusters displaying dose effects as a function of dietary sodium nitrite content in D. Clusters are agglomerated at the genus level and normalized Log_10_ abundances were represented using scatter plots with bar (mean ± sem). * p ≤ 0.05; **p ≤ 0.01.

#### 3.1.5. Sodium nitrite reduction or removal decreased promotion of colon carcinogenesis in the rat model

Compared to the cooked ham model produced with 120 mg/kg of sodium nitrite (Ni-120), the reduction of sodium nitrite to 90 mg/kg (Ni-90) induced a reduction in the number of the preneoplastic lesions MDF per colon, while the removal (Ni-0) induced an effect very close to significance (p = 0.069) (Figure 4A). Nitrite salt reduction and removal induced a clear and significant reduction of the total number of crypts depleted in mucin (Figure 4A and B). Globally the same reducing effect was observed considering only bigger MDF lesions (Figure 4B), with, a slightly less significant effect. However, nitrite salt removal that is protective against processed meat-induced promotion of carcinogenesis in comparison to Ni-120 was not more effective than Ni-90.

**Figure 4.**
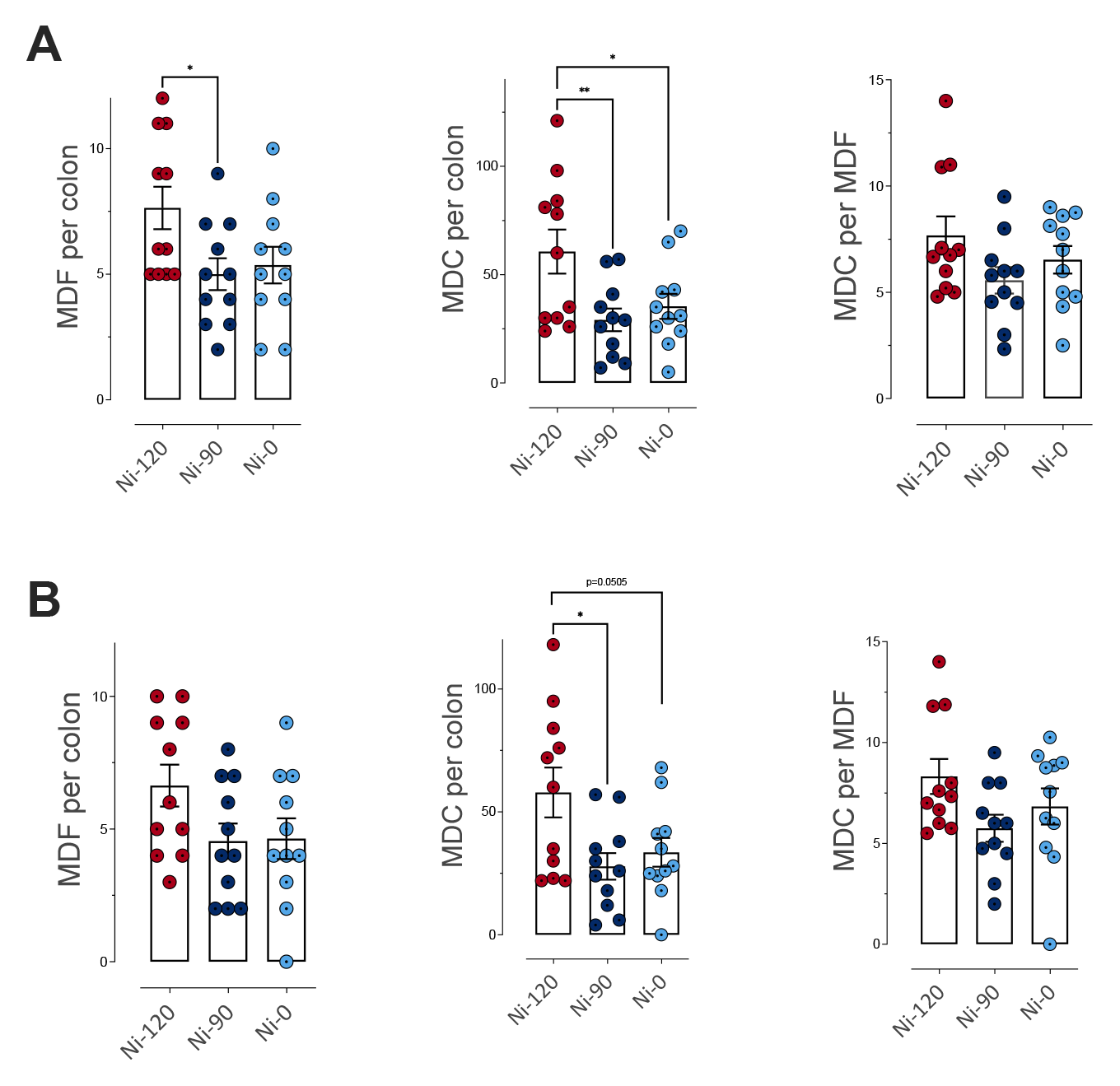
Impact of sodium nitrite levels in cooked ham models (0 *vs* 90 *vs* 120 mg/kg) on MDF formation in rat colon. **A**-Number of MDF per colon, of mucin depleted crypts (MDC) per colon and crypts per focus for MDF with a multiplicity (i.e. the number of crypts forming each focus) higher than 2 crypts/MDF. **B**- Number of MDF per colon, of MDC per colon and crypts per focus for MDF with a multiplicity higher than 4 crypts/MDF. Data were represented using scatter plots with bar (mean ± sem), * p ≤ 0.05; **p ≤ 0.01.

### 3.2. Impacts of three alternatives to nitrite salt on cooked ham model products, microbiological risk, colon ecosystem and colorectal carcinogenesis

#### 3.2.1. Consequences of use of three alternatives to nitrites on nitrosylation, nitrosation and lipid peroxidation in the cooked ham model

In comparison to the reference cooked ham model with 120 mg/mg of sodium nitrite (Ni-120), alternatives to sodium nitrite salt had very different effects on the characteristics of processed meats (Table 2). The use of a vegetable stock (VS) induced an increase in the percentage of heme iron nitrosylation, the concentration of residual nitrite compared to the reference processed meat Ni-120 (p<0.05). The VS alternative had no impact on the concentration of total non-volatile *N*-nitrosamines, lipid peroxidation or residual nitrates, while it decreased the concentration of nitrosothiols in comparison to Ni-120 (Table 2). The Polyphenol-Rich Extract (PRE) increased the concentration of total non-volatile *N*-nitrosamines (p ≤ 0.05, Table 2) and decreased significantly the concentration of nitrosothiols, residual nitrites and nitrates without impact on lipid peroxidation in comparison to Ni-120. The processed meat produced with the Lallemand solution (YE) presented significantly less nitrosylated heme iron, total non-volatile *N*-nitrosamines, nitrosothiols and residual nitrates or nitrites and a strong and significant increase of lipid peroxidation (TBARS) in comparison to the reference processed meat Ni-120 (Table 2).

**Table 2:** Impact of alternatives to nitrites on biochemical characteristics of cooked ham models. Impact of 120 mg/kg of sodium nitrite or Vegetable Stock (VS), Polyphenol-rich Extract (PRE), Lallemand solution (YE). Data were mean ± SD). Mean values with unlike letters were significantly different (p ≤ 0.05). NA: data not available due to an interference with polyphenols contained in PRE (see supplementary data (Table S6 and S7)

|  | Ni-120 | VS | PRE | YE |
| --- | --- | --- | --- | --- |
| **Total heme iron ($\mu\text{M}$)** | 189 ± 40 <sup>$\alpha$</sup> | 153 ± 12 <sup>$\alpha$</sup> | 156 ± 18 <sup>$\alpha$</sup> | 142 ± 2 <sup>$\alpha$</sup> |
| **Nitrosylated iron ($\mu\text{M}$)** | 110 ± 11 <sup>$\alpha$</sup> | 108 ± 7 <sup>$\alpha$</sup> | NA | 20 ± 2 <sup>$b$</sup> |
| **% Nitrosylated iron** | 58 | 70 | NA | 14 |
| **TBARS ($\mu\text{g MDA/g ham}$)** | 0.041 ± 0.006 <sup>$\alpha$</sup> | 0.032 ± 0.002 <sup>$\alpha$</sup> | 0.041 ± 0.001 <sup>$\alpha$</sup> | 0.158 ± 0.021 <sup>$b$</sup> |
| **Total free iron (mg/kg)** | 4.92 ± 0.04 <sup>$\alpha$</sup> | 4.21 ± 0.01 <sup>$b$</sup> | 2.42 ± 0.01 <sup>$c$</sup> | 5.48 ± 0.03 <sup>$d$</sup> |
| **Total non volatile *N*-nitrosamines (mg/kg)** | 2.09 ± 1.02 <sup>$\alpha$</sup> | 1.57 ± 0.88 <sup>$\alpha$</sup> | 7.09 ± 2.24 <sup>$b$</sup> | 0.55 ± 0.14 <sup>$c$</sup> |
| **Nitrosothiols (mg/kg)** | 2.05 ± 0.25 <sup>$\alpha$</sup> | 0.68 ± 0.47 <sup>$b$</sup> | 0.98 ± 0.33 <sup>$b$</sup> | 0.15 ± 0.03 <sup>$c$</sup> |
| **Residual nitrites (mg/kg)** | 9.36 ± 0.23 <sup>$\alpha$</sup> | 13.56 ± 0.50 <sup>$b$</sup> | 0.19 ± 0.03 <sup>$c$</sup> | 0.08 ± 0.02 <sup>$c$</sup> |
| **Residual nitrates (mg/kg)** | 24.95 ± 1.07 <sup>$\alpha$</sup> | 23.35 ± 1.04 <sup>$b$</sup> | 2.79 ± 1.03 <sup>$c$</sup> | 3.20 ± 0.10 <sup>$c$</sup> |

#### 3.2.2. *Listeria monocytogenes* growth in sliced cooked ham model products as function of alternatives to nitrites

After a week of storage, *L. monocytogenes* was not able to grow on sliced cooked ham model products, regardless of the recipes (δ < 0.5 Log_10_ CFU/g, Figure 5). Thereafter, all products supported the growth of the pathogen (δ > 0.5 Log_10_ CFU/g) until the end of storage. However, the products with polyphenol-rich extract (PRE) or Lallemand solution (YE) exhibited significantly higher growth potentials than those manufactured with the vegetables stock (VS) or with an ingoing amount of 120 mg/kg of sodium nitrite from day 14 to day 22 (p ≤ 0.0001, Figure 5).

**Figure 5.**
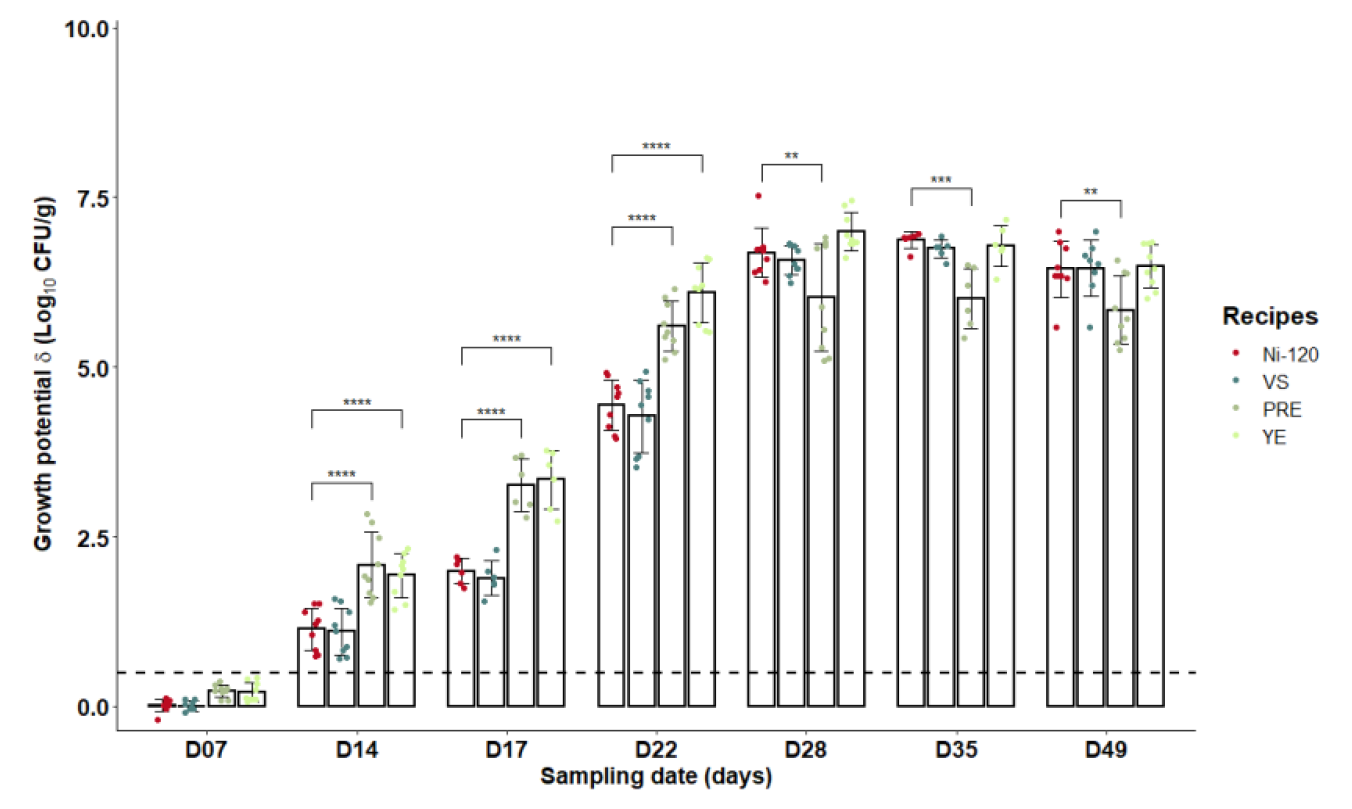
Growth potentials (δ) of *Listeria monocytogenes* at several sampling times during shelf-life of the sliced cooked ham model products with alternatives to nitrites. (120 mg/kg of nitrite or Vegetable Stock (VS), Polyphenol-rich Extract (PRE), Lallemand solution (YE)). Data were represented using scatter plots with bar (mean ± SD); **p < 0.01, ***p < 0.001, ****p < 0.0001. The dashed line represents the limit value of 0.5 Log_10_ CFU/g above which the cooked ham samples support the growth of *L. monocytogenes*.

#### 3.2.3. Comparison of the impacts on endogenous formation of nitroso compounds, luminal and global peroxidation or cytotoxic and genotoxic fecal activities of nitrites or alternatives tested

As observed for the biochemical characteristics of cooked ham models, the three alternatives to sodium nitrite have different effects on endogenous reactions. In comparison to Ni-120, the use of vegetable stock (VS) induced a significant increase of the fecal endogenous formation of total nitroso compounds and nitrosyl iron (p ≤ 0.05, Figure 6A) without altering the production of fecal RNNO and RSNO (Figure 6A), luminal lipid peroxidation, urinary excretion of DHN-MA (Figure 6B) or heme bioavailability (Figure 6C). The replacement of sodium nitrite by polyphenol-rich extract (PRE) induced a significant increase of fecal endogenous formation of total nitroso compounds, nitrosyl iron and RSNO (p < 0.05, Figure 6A) while strongly and significantly reducing RNNO level (p ≤ 0.05, Figure 6A). The consumption of PRE cooked ham model did not change luminal lipid peroxidation (TBARS) or heme bioavailability (Figure 6C), but decreased significantly urinary excretion of DHN-MA (Figure 6B).

**Figure 6.**
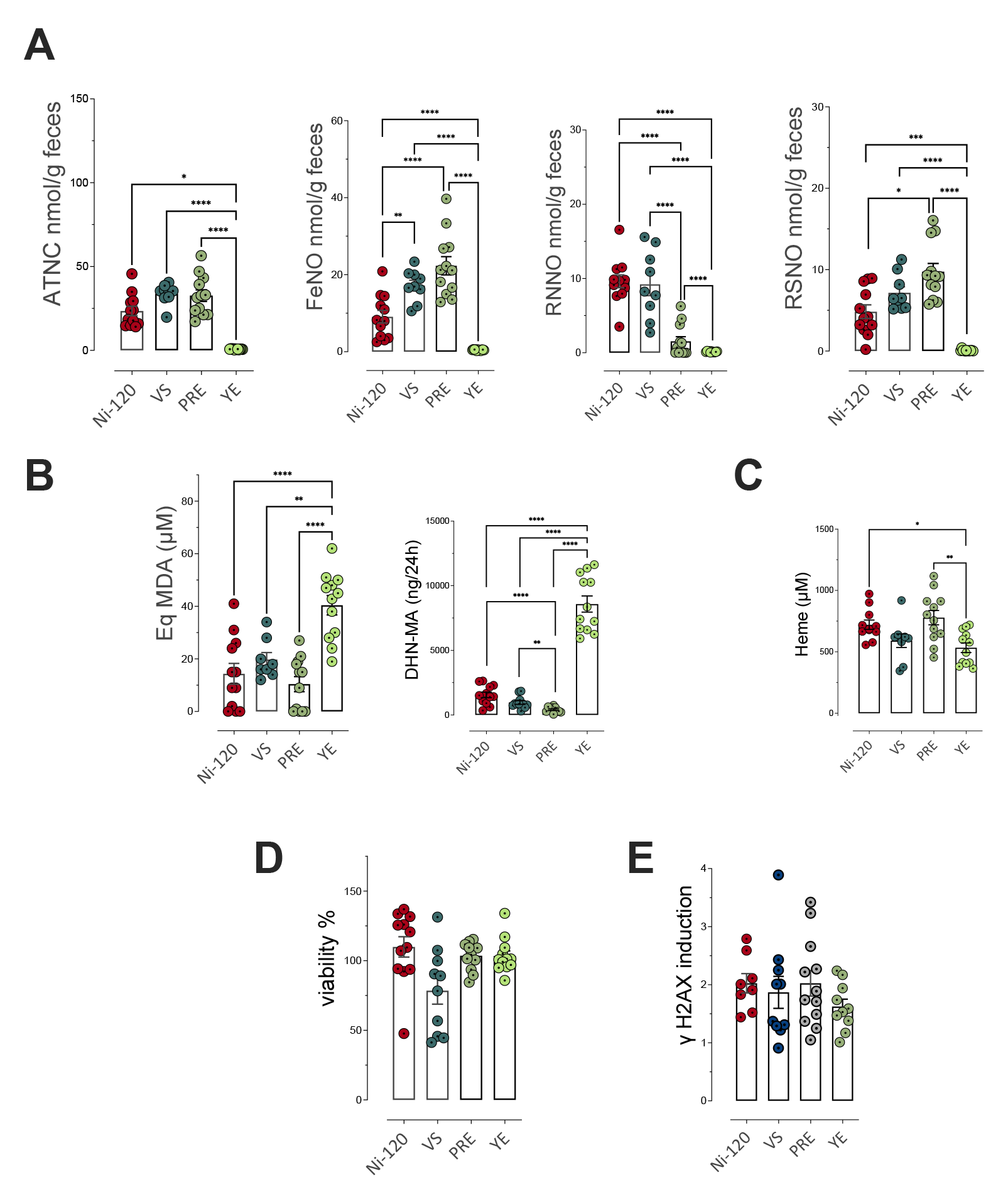
Impact of nitrite salt alternatives (Vegetable Stock (VS), Polyphenol-rich Extract (PRE), Lallemand solution (YE)) on fecal and urinary biomarkers of NOCs formation and lipid peroxidation. **A**-Nitroso-compounds in fecal water measured as total NOCs (ATNC), as nitrosyl iron (FeNO), *N*-nitroso compounds (RNNO) and S-nitrosothiols (RSNO) (nmol/g of feces). **B**-Lipid peroxidation measured as TBARS (MDA equivalents, µM) in fecal water and DHN-MA in urine of 24h (ng/vol of 24h). **C**-Heme in fecal water (µM). **D**- Cytotoxic activity of fecal water as % of cellular viability. **E**- Genotoxic activity of fecal water. Data were represented using scatter plots with bar (mean ± sem), * p ≤ 0.05; **p ≤ 0.01.

With an opposite effect, the cooked ham model treated with the Lallemand solution (YE) induced a strong and significant decrease of fecal endogenous formation of total nitroso compounds, nitrosyl iron, RNNO and RSNO (p < 0.05, Figure 6A), but a high increase of luminal lipid peroxidation, and particularly of urinary excretion of DHN-MA (Figure 6B) without modification of the heme bioavailability comparing to sodium nitrite use (Figure 6C). These modifications in the fecal contents of *N*-nitroso compounds and alkenals were not associated with a change in the cytotoxic and genotoxic activities of fecal waters (Figure 6D & E).

#### 3.2.4. Effect of nitrite alternatives on community distribution and diversity of the fecal microbiota as determined by 16S rRNA gene sequencing

As compared with distribution at the order level within microbiota of rats Ni-120 (Figure 7A), changes in response to sodium nitrite alternatives were observed within *Bifidobacteriales* (increase with YE only, Figure S7A), *Desulfovibrionales* (decrease mainly with YE), *Peptococcales* (increase with VS and PRE, Figure S7) and *Peptostreptococcales-Tissierellales* (decrease with PRE and YE, Figure S7). Regarding diversity, none of the indices related to *α*- diversity were altered by diet content (Figure 7B), but dissimilarity analysis between groups (p = 0.0004, Figure 7C) indicated that the difference in microbiota between Ni-120 and YE was significantly greater (p = 0.001) than that between Ni-120 and PRE (p = 0.017), whereas no significant difference was obtained between the microbiota of the Ni-120 and VS rat groups (p = 0.083). Differential analysis at the genus level supported these results (Figure 7D, Table S6B) and revealed distinct signatures depending on the alternative tested (Figure 7E, F, G, Figure S7). Interestingly, abundance variations for some genera in response to diet supplemented with VS (Figure 7E), PRE (Figure 7F) or YE (Figure 7G) were similar to those observed in diets in which sodium nitrite content was reduced or removed (Figure S7B).

**Figure 7.**
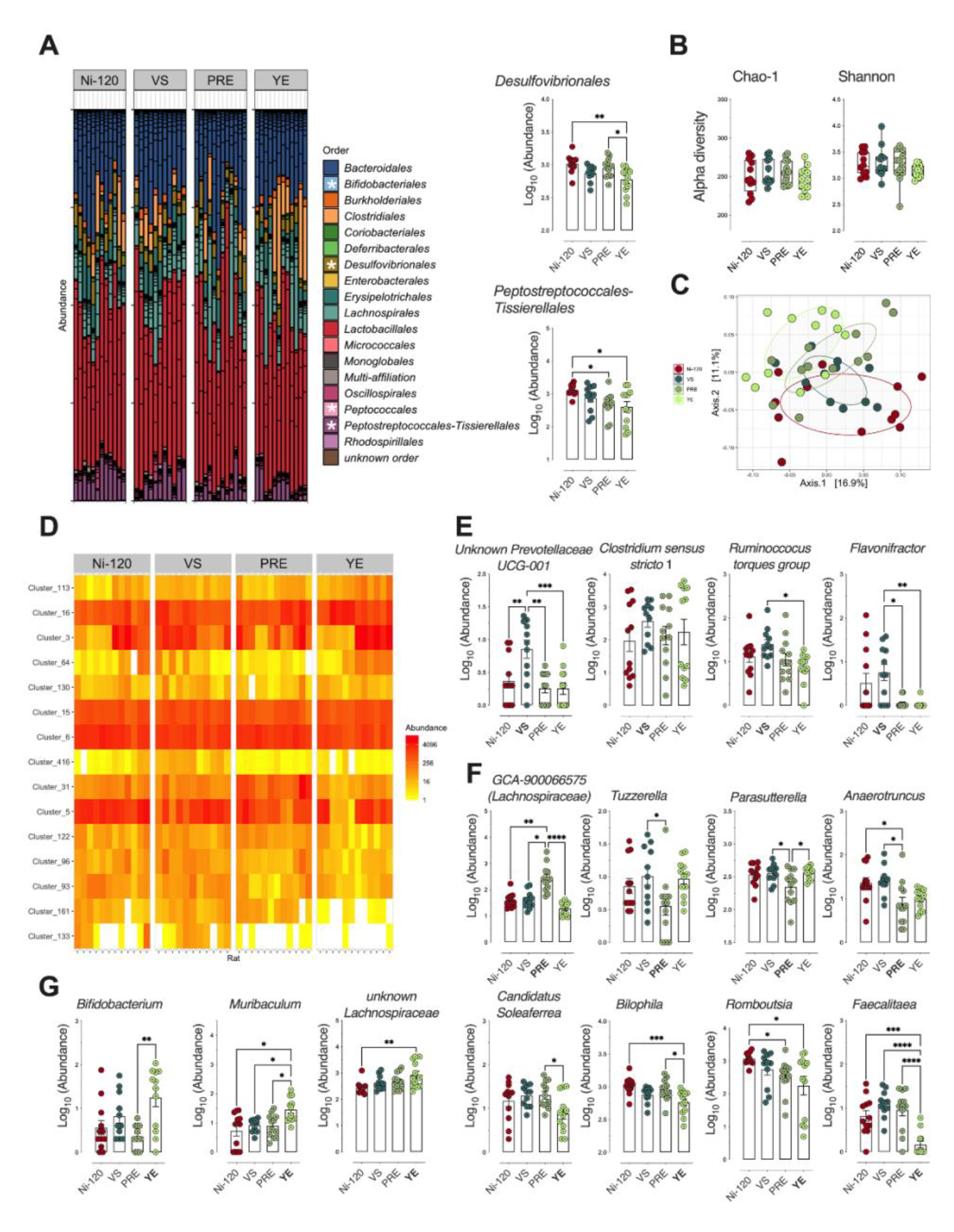
Impact of alternatives to sodium nitrites 120 mg/kg (Vegetable Stock (VS), Polyphenol-rich Extract (PRE), Lallemand solution (YE) on fecal microbiota. **A**-Distribution of bacterial communities at the order level. *Main impact on *Bifidobacteriales*, *Desulfovibrionales*, *Peptococccales* and *Peptostreptococcales-Tissierellales* using differential abundance analysis at the order level (Deseq2, *P*_adj_≤0.05): Normalized Log_10_ abundances were represented using scatter plots with bar (mean ± sem). **B**-Alpha diversity: No significant impact seen on richness (Chao-1) or eveness (Shannon). Individual values are represented using box and whiskers (+ mean). **C**-Beta diversity (Unifrac distances, manova P≤0.001). The rats fed the 4 meat-based diets clustered differentially according to their respective microbiota in terms of qualitative abundance and taxonomy of OTUs. **D**-Heatmap of clusters agglomerated at the genus level affected by the meat-based diets (15 clusters). *P*_adj_≤0.05 using differential abundance analysis (Deseq2). **E**-Normalized abundance of the 4 clusters in D displaying specificities associated with the fermented Vegetable Stock diet (**VS**). Clusters are agglomerated at the genus level and normalized Log_10_ abundances were represented using scatter plots with bar (mean ± sem) **F**-Normalized abundance of the 4 clusters in D displaying specificities associated with the Polyphenol-rich Extract diet (**PRE**). Clusters are agglomerated at the genus level and normalized Log_10_ abundances were represented using scatter plots with bar (mean ± sem). **G**-Normalized abundance of the 7 clusters in D displaying specificities associated with the Lallemand solution (**YE**). Clusters are agglomerated at the genus level and normalized Log_10_ abundances were represented using scatter plots with bar (mean ± sem). * p ≤ 0.05; **p ≤ 0.01; *** p ≤ 0.001.

#### 3.2.5. Lack or low impact of alternatives to nitrites tested on promotion of colon carcinogenesis

Compared to the cooked ham model produced with 120 mg/kg of sodium nitrite (Ni-120), the various alternatives did not modify significantly the number of preneoplastic lesions (Figure 8 A and B), despite a decreasing trend for VS and PRE compared to Ni-120. If considering the number of mucin depleted crypts, both ANOVA reveal an almost significant effect (p = 0.064 and p = 0.071), for all lesions and for lesions with 4 or more crypts, respectively.

**Figure 8.**
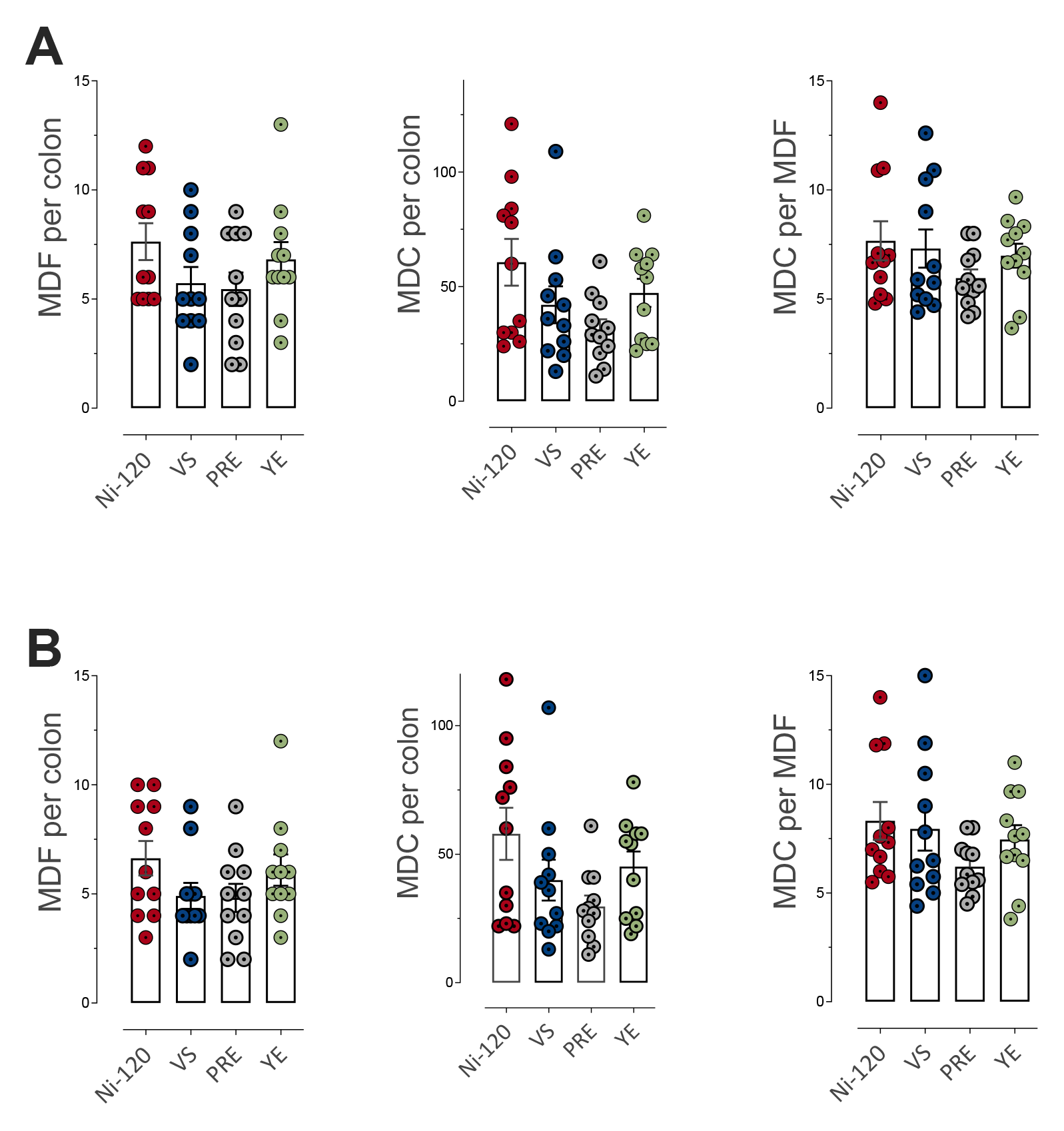
Impact of alternatives to sodium nitrite (Vegetable Stock (VS), Polyphenol-rich Extract (PRE), Lallemand solution (YE)) on MDF formation in rat colon. **A**-Number of MDF per colon, of mucin depleted crypts (MDC) per colon and crypts per focus for MDF with a multiplicity (i.e. the number of crypts forming each focus) higher than 2 crypts/MDF. **B**- Number of MDF per colon, of MDC per colon and crypts per focus for MDF with a multiplicity higher than 4 crypts/MDF. Data were represented using scatter plots with bar (mean ± sem).

## 4. Discussion

This study conducted for the first time, to the best of our knowledge, a multi-risk assessment of the consequences of the reduction, removal or substitution of sodium nitrite in a cooked ham model, regarding impacts on (i) growth potential of *L. monocytogenes*, (ii) the meat product characterization, (iii) the promotion of colorectal carcinogenesis (at a preneoplastic stage) and (iv) the colonic ecosystem (endogenous neoformations of lipid peroxidation products and NOCs, and microbiota dysbiosis).

Our results clearly showed that nitrite salt reduction or removal in cooked model ham could modify most of the tested parameters in this work (Table 1, Figures 1-4). As observed previously ^[21]^, the results of the present experimentation confirmed that nitrite salt removal limited colorectal carcinogenesis promotion at the preneoplastic stage (Figure 4). MDF, used in these studies to assess the impact on promotion of colorectal carcinogenesis, were identified in 2003 by Caderni *et al.*^[26]^ in a CRC animal model and in humans in 2008 by the same group.^[38]^ MDF share dysplastic traits and mutational profiles with colonic tumors as β- catenin and frequent Apc mutations.^[39]^ MDF are proposed as cancer precursors and usable endpoints in short term carcinogenesis study with a good predictive character for tumor stage effects.^[26]^ In this way, the very recent results of Crowe *et al.*^[40]^ confirmed our previous results at the tumor stage, by demonstrating also in *Min* mice that a nitrite-free sausage protects against promotion of intestinal tumors in comparison to nitrited ones. The results of our team, including the present study, and the results of Crowe *et al.* support the role of nitrite salt food additive in colorectal cancer promotion associated to processed meat consumption. These data are also in agreement with the positive epidemiological associations between the intake of the nitrited and nitrated food additives and the development of cancers in the NutriNet-Santé French cohort, including a positive trend for colorectal cancer.^[20]^

Evaluated for the first time in the present study in an animal model of colorectal carcinogenesis, reduction of sodium nitrite from 120 to 90 mg/kg also induced a protective effect on colon preneoplastic lesions (Figure 4). But, importantly, if the number of preneoplastic lesions was also decreased with the removal, we did not observe any difference between reduction and removal. As expected, residual nitrite and nitrate salts were dose-dependently decreased in the cooked ham models with the three levels of nitrites, while the total nitroso compounds and the three subcategories were found decreased in the same way in the feces of the rats fed the ham-rich diets (Figure 2). However, the effect of this reduction or removal of sodium nitrite remains small on colon fecal microbiota, as revealed by the lack of structural difference in α and β diversity (Figure 3B-C) as well as on gene expressions in colon mucosa (Figure S4), urinary excretion of 8-isoPGF_2α_ (Figure S6A) fecal cytotoxic and genotoxic activities (Figure 2D-E). Regarding fecal microbiota at the order level, we observed a decrease in *Peptostreptococcales-Tissierellales* (Figure 3A), some members of which, such as *Peptostreptococcus*, are known to be enriched in fecal and mucosal samples from patients with CRC^[41]^. In our rat model, the decrease in *Peptostreptococcales-Tissierellales* was mainly due to the decrease in *Romboutsia* (Figure 3D-E), the presence of which has found associated in mice with intestinal damage, increase of inflammatory cytokines, iNOS and AOM-DSS-induced CRC.^[42][43]^ But, as a consequence of nitrite salt removal, and as already reported by our group^[21]^, lipid peroxidation as measured by the TBARS index was increased in the cooked ham model (Table 1). The TBARS index in rat feces also was significantly increased when nitrite salt was removed from the cooked ham model, while the urinary lipid peroxidation biomarker DHN-MA was increased dose-dependently as nitrite salt decreased (Figure 2B). Numerous studies of our team have proposed experimentally a role for lipid peroxidation products in the promotion of colorectal carcinogenesis ^[23,44,45]^ Thus, despite the absence of fecal nitrosation and nitrosylation when sodium nitrite was removed (Figure 2A), the lack of additional protection against cancer risk in comparison to the decrease at 90 mg/kg could be explained by this significant and concomitant increase in luminal lipid peroxidation. Moreover, the high level of DHN-MA observed in the present study reflected a significant intestinal absorption of those toxic alkenals and raised the question of a systemic impact. Indeed, the studies of Awada *et al.*^[46]^ and Bolea *et al.*^[22]^ demonstrated that an increase in luminal alkenal production is associated, as reflected in our study by the increase of the fecal TBARS index, to an increase in plasma concentrations of bioactive alkenals. The study of Bolea *et al.* also reported in mice that a high plasmatic level of alkenals caused dysfunction in peripheral tissues, particularly vascular dysfunction.

However, previous results from our team have demonstrated in a CRC animal model that enriching a cooked processed meat with antioxidants, such as vitamin E, red wine or pomegranate extracts, limited the processed meat-induced luminal lipid peroxidation and DHN-MA urinary excretion.^[24,47]^ The efficacy of vitamin E supplementation has been verified also in healthy human volunteers.^[47]^ Thus, on the basis of all these data, we propose that the protective efficiency of sodium nitrite removal on colon carcinogenesis could be improved by controlling luminal lipid peroxidation with natural antioxidants. Furthermore, by limiting the luminal formation of alkenals, their absorption will be limited and their effects on peripheral tissues should therefore be also controlled. The decrease in DHN-MA urinary excretion after processed meat supplementation with red wine or pomegranate extracts supports this hypothesis.^[24]^ Future studies should focus on these points.

Regarding the challenge test assays on *L. monocytogenes*, cooked ham model samples used in the microbiological assays exhibited typical features (more details in supplementary material and Table S4 and S5) and the nitrate, nitrite and salt (NaCl) levels measured were consistent with the additive doses in corresponding recipes (see Table S3-S4). The present study showed that sodium nitrite added in the cooked ham products exerted an inhibitory effect against *L. monocytogenes*, as previously reported on various ready-to-eat cooked meat products.^[15–17,48]^ Although sodium nitrite alone was not sufficient to prevent the growth of *L. monocytogenes* during actual shelf life of sliced cooked ham products, ingoing amounts of sodium nitrite equal or greater than 90 mg NaNO_2_ / kg led to a significant reduction of *L. monocytogenes* growth during the first 3 weeks of storage compared to the control without nitrite (Ni-0). Moreover, it should be noted that the presence of sodium ascorbate (0.3 g/kg) in the different recipes (Ni-120, Ni-90) may have enhanced the anti listerial effect of sodium nitrite as previously described by others.^[15,48]^

Among the three alternative products that were tested as nitrite salt replacement, the vegetable stock (VS) gave, for almost all the parameters tested, results that were very close to those of Ni-120, whatever the dose of the food additive. Indeed, results on luminal lipid peroxidation and ATNC formation with different concentrations of vegetable stock are similar to nitrites doses of 120, (Figure 6A-B), 80 or 40 mg/kg (Figure S1A-B). The yeast/bacterial extract (YE) was unable to control lipid peroxidation in the processed meat product (Table 2) and *in vivo* (Figure 6B). As such, this alternative can be compared to nitrite removal (Ni-0), with however a possible slight gain in the modulation of some microbiota specific species: regarding the commensal microbiota, YE helped to improve microbiota homeostasis by increasing the following bacterial genus (*Bifidobacteria*, *Muribaculum* and *unknown Lachnospiraceae* associated respectively with anti-cancer effects^[49]^, longevity ^[50]^ and short-chain fatty acid production^[51]^) and decreasing genus associated with adverse effects (*Bilophila, Romboutsia* and *Faecalitaea* associated respectively with inflammation ^[52]^, intestinal damage, increase of inflammatory cytokines ^[42]^ and presence of adenoma in patients ^[53]^). However, these modifications were not associated with a significant health positive effect on preneoplastic lesions and were correlated with very low impacts on gene expressions in colon mucosa (Figure S5), urinary excretion of 8-isoPGF_2α_ (Figure S6B) and fecal cytotoxic and genotoxic activities (Figure 6D-E). Ni-0 and YE formulations do not provide nitrosated compounds and do not improve the health effect that could be expected in comparison to nitrite reduction (Ni-90), most likely because of the lack of lipid peroxidation control. This is even more obvious for YE, in which there is no ascorbic acid, and this absence can explain an even greater increase in lipid peroxidation. These results underline the fact that it seems very important to control this process. The addition of antioxidants could represent a solution and the PRE product could provide a valuable alternative. However, the presence of a little concentration of non-volatile nitrosamines in the PRE-treated ham model and of some ATNC and nitrosyl iron in feces, makes us think that there is maybe a source of NO in this product. This was not expected as this curing product is not supposed to provide any NO source. It was not possible to determine unequivocally FeNO in PRE cooked ham model by the colorimetric method we used because interferences due to zinc protoporphyrin (ZnPP), known for giving Parm ham its pink/red colour, or to the presence of colored polyphenols cannot be excluded. However, Zn quantification in the nitrited (Ni-120) and PRE-treated ham did not reveal any difference (Table S1). The percentage of FeNO detected increased when loading Ni-0 ham with the polyphenol epigallocathechine gallate known to bring a pink colour (Table S6). This increase is even amplified when adding ferulic acid and ascorbic acid (FEA) (Table S7). Interestingly, the determination of NO_2_, NO_3_, RSNO and RNNO using the colorimetric Griess method did not interfere with FEA (Table S8). Nonetheless, concerning ATNC and FeNo detection in feces, the method we used is a published method that has been widely used and usually gives consistent results, to the best of our experience. This assay method has been used, for example, by the group of S. Bingham and G. Kuhnle, who are references in the field of endogenous formation of NOCs during consumption of meat products.^[54,55]^ The CLD88 assay has also been used by this team to monitor the fecal excretion of NOCs in humans after consumption of processed red meat with standard or reduced levels of nitrite and added polyphenols.^[56]^ However, this method remains undirect (detection of NO by chemiluminescence) and a possible interference due to components in the PRE product cannot be fully excluded. To explore this interference hypothesis, we overloaded Ni-0 fecal water with polyphenols and ascorbic acid and analyzed the ATNC before and after this overloading (Fig S3). No differences were observed, indicating the absence of interference and therefore of false positives attributable to polyphenols in the PRE formulation. Finally, the presence of non-volatile nitrosamines in the PRE cooked ham model, although in low concentration, led us to suspect a nitrosating and nitrosylating agent in this recipe, with thus a consequent risk on the formation of volatile nitrosamines considered carcinogenic. However, the determination of 5 volatile nitrosamines in the reference cooked ham model (Ni-120) and the three alternatives showed values below the quantification limit of these compounds (Table S2).

Regarding the challenge test assays on *L. monocytogenes*, the use of the vegetable stock (VS) led to a similar reduction of *L. monocytogenes* growth during the first 3 weeks of storage compared to that obtained with an ingoing amount of 120 mg/kg of sodium nitrite (Ni-120). The inhibitory effect of the VS alternative (with high sodium nitrate content) against *L. monocytogenes* is not surprising since natural nitrate is reduced to nitrite (to reach an expected concentration of 80 mg NaNO_2_/kg in our assays) during food processing *via* the nitrate reductase activity of starter cultures.^[57]^ Alternatively, certain plant active extracts were demonstrated to limit the growth of foodborne pathogens including *L. monocytogenes* on that kind of product, which is often attributed to high contents of polyphenolic compounds.^[13,58–61]^ However, in our assays, the alternative PRE employed following the manufacturer’s recommended concentration was less efficient than sodium nitrite (≥ 90 mg/kg) or the VS alternative solution in reducing *L. monocytogenes* growth during the first 3 weeks of storage. A similar trend was obtained using the newly developed strategy Lallemand solution (YE) to replace or reduce nitrite salts.

Based on this study co-evaluating for the first time the effects of reduction, removal and alternatives to sodium nitrite, the decrease of sodium nitrite salt level in cooked processed meat appears to be a short-term perspective for limiting population exposure to sodium nitrite additive, to nitrosated compounds and to colon carcinogenesis risk while slowing the *L. monocytogenes* growth and efficiently controlling lipid peroxidation. The lack of a greater effect on carcinogenesis risk with removal highlights the need to consider the increase in luminal lipid peroxidation. This conclusion is reinforced by the lack of protective effect of the YE which, if it is truly free of nitrites/nitrates or NO, induced a very high luminal lipid peroxidation.

In conclusion, if the removal of nitrite salts should not be excluded in certain cooked meats, in particular those with a very short shelf life, it appears necessary to quickly evaluate if the enrichment with antioxidants of low-nitrited or non-nitrited cooked processed meat (like the cooked ham model of this study) would allow a greater protection than with the reduction to 90 ppm. This work also highlights the need for an evaluation of alternatives on endogenous formations and colon carcinogenesis promotion before introduction on the market.

## Supporting Information

Supporting Information is available from the author.

## Abbreviations

ATNC: apparent total nitroso compounds
CRC: colorectal cancer
DHN-MA: 1,4-Dihydroxynonane mercapturic acid
DSS: dextran sodium sulfate
FeNO: nitrosyl iron
FW: fecal water
HNE: 4-hydroxynonenal
MDF: mucin depleted foci
NOCs: nitroso-compounds
RNNO: residual *N*-nitroso compounds
RSNO: *S*-nitrosothiols
TBARS: thiobarbituric acid reactive substances.

## Acknowledgments

The authors thank Xavier Blanc (UE 1298 SAAJ, Sciences de l’Animal & de l’Aliment, INRAE) for providing custom experimental diets and all members of the EZOP (Animal facility) for assistance with the animal experimentation. The authors thank the Genotoul bioinformatics platform Toulouse Occitanie and Sigenae group for providing help and storage resources thanks to Galaxy instance (https://galaxy-workbench.toulouse.inra.fr). The authors thank Bettina Seeger (University of Veterinary Medicine, Hannover) for providing the CLD88 et NO-Analyzer for NOC quantification. The authors thank G. Kuhnle for fecal NOC analyses in the supplementary experiment.

## Conflict of Interest

A P Promeyrat, B F, J-L M Martin, S Jeuge and G Nassy were employed by the French Pork Institute (IFIP). This study was co-financed (50%) by IFIP. F Pierre, F Guéraud and V Santé-Lhoutellier have some research projects from their academic research teams that have been co-financed by the processed meat sector.

## Author Contributions

Françoise Guéraud, Fabrice Pierre, Véronique Santé-Lhoutellier, Gilles Nassy and Aurélie Promeyrat designed the research studies, analyzed data, and wrote the manuscript. Bastien Frémaux, Maïwenn Olier acquired and analyzed data and wrote the manuscript. Tina Kostka and Giovanna Caderni acquired data, analyzed data, reviewed and edited the manuscript. Charline Buisson, Nathalie Naud, Edwin Fouché, Jacques Dupuy, Eléna Keuleyan, Noémie Petit, Valérie Bézirard, Laurent Aubry and Jean-Luc Martin conducted experiments, acquired and analyzed data. Sabine Jeuge, analyzed data. Vassilia Théodorou, Pascale Plaisancié, Cécile Héliès-Toussaint reviewed and edited the manuscript.

## Data Availability Statement

The sequencing data of the 16S rRNA gene have been deposited in the European Nucleotide Archive (ENA) at EMBL-EBI under accession number PRJEB59736 (https://www.ebi.ac.uk/ena/browser/view/PRJEB59736).

