## Supplementary material for "Reduction, removal or replacement of sodium nitrite in a model of cured and cooked meat: a joint evaluation of consequences on microbiological issues in food safety, colon ecosystem and colorectal carcinogenesis"

-------

-------

#### 1. Supplementary experiment: impact of sodium nitrite concentrations and vegetable stock in a cooked ham model on fecal and urinary biomarkers of lipid peroxidation and on NOC formation

##### *1.1 Supplementary Experimental Section*

*-Processed meat production and composition.*

For this short-term nutritional study, 4 experimental cooked ham models were made in a pilot-scale unit (IFIP, France) according to a process that permitted a homogeneous curing treatment. These products were composed of 2.5 kg ground pork supplemented with a specific brine solution as described below. They were prepared with fresh boned and derinded pork ham. Further trimming and cutting were performed, and the pieces of meat were ground through a 20 mm plate of a grinder (DRC 98, PSV, France). In each experiment, the ground pork was divided into 4 recipes, then mixed with appropriate concentrations of sodium nitrite (80, or 40 mg/kg) or alternative (vegetable stock with Nitrated broth NAT 223 (Soussana , France) and Starters CS300 (Soussana, France) to mimic 80 or 40 mg/kg of sodium nitrite in VS80 and VS40 with respectively 4 and 2 g of broth per kg of meat and 0.25 g of starter by kg of meat), NaCl (14 g/kg), sodium erythorbate (500 mg/kg), dextrose (5 g/kg) using a vacuum mixer (CDH, France) at a constant speed of 350 rpm for 10 min. The starter used in the VS recipes was stored at -20°C and prepared according to the manufacturer's instructions. In all recipes, water was added in order to have a 10% brine rate.

After processing, the 4 experimental cooked ham models were stored at -20°C until the nutritional study in rats.

*-Nutritional study (14 day-long):*

*Experimental design:* Male Fischer 344 (F344/IcoCrl) rats (5 rats/group) were purchased from Charles River Laboratories at 5 weeks of age. After acclimatization, the animals were randomly assigned to the experimental groups and given diets for two weeks. Rats were then euthanized.

Feces were collected in plastic metabolic cages respectively at days 7-9 and 14 and urines were collected at days 7 and 14 and to follow fecal and urinary biomarkers of peroxidation, nitrosylation and nitrosation.

*Experimental diets:* Experimental diets were based on powdered low-calcium, no-fat AIN-76 rodent diets (SAAJ, Jouy-en-Josas, France), supplemented with 5% safflower oil (MPBio) given in a separated feeder (7.5 g/d/rat) and 30 g of processed meat produced with nitrites at 40 or 80 mg/kg (respectively Ni-40 and Ni-80) or vegetable stock mimicking same sodium nitrite level (respectively VS-40 and VS-80) corresponding to 46-49% of dry matter of the diet.

*Analysis of thiobarbituric acid reactive substances (TBARS), heme iron, ATNC and nitrosyl iron in feces, and 1,4-dihydroxynonane mercapturic acid (DHN-MA)in urine.*

TBARS, heme iron and DHN-MA: see *Experimental Section* of the main article except for the heme assay : heme iron was measured by fluorescence in fecal water according to Sesink *et al.* as already described[1].

ATNC analysis: ATNC were analyzed using a method slightly different from the one used in the main manuscript [2], using a CLD88 Exhalyzer (Ecomedics, Duernten, Switzerland). Sulfamic acid solution (500 µl, 5%) was added to 100 µl of fecal water to remove nitrite and samples were injected into a purged vessel kept at 60°C and filled with a standard tri-iodide reagent (38 mg $I_2$ was added to a solution of 108 mg KI in 1 mL water; to this mixture, 13.5 mL glacial acetic acid was added) to determine total ATNC. To determine mercury(II) stable compounds, 100 µL of 10 mM aqueous $HgCl_2$ was added prior to analysis; to determine mercury(II) and ferricyanide stable compounds, 100 µL each of 10 mM aqueous $HgCl_2$ and 10 mM aqueous $K_3Fe(CN)_6$ solution were added prior to analysis. Nitrosyl iron was determined as difference between mercury(II) stable ATNC and mercury(II) and $K_3Fe(CN)_6$ stable compounds. Data are concentrations (in µM), measured in triplicate in 100 µL of each sample.

##### *1.2 Supplementary Result Section*

The consumption of processed meats did not induce a significant difference between the 4 experimental groups (see Figure S1). The presence of sodium nitrite (40 or 80 mg/kg of meat) or vegetable stock (mimicking the intake of 40 or 80 mg of nitrites/kg of meat) in processed meats did not modify fecal (Figure S1A/TBARS) and global (Figure S1A/DHN-MA) lipid peroxidation biomarkers, the formation of total nitroso compounds (Figure S1B/ATNC) and nitrosylated iron (Figure S1B/FeNO) as well as fecal heme iron bioavailability (Figure S1C) in rats.

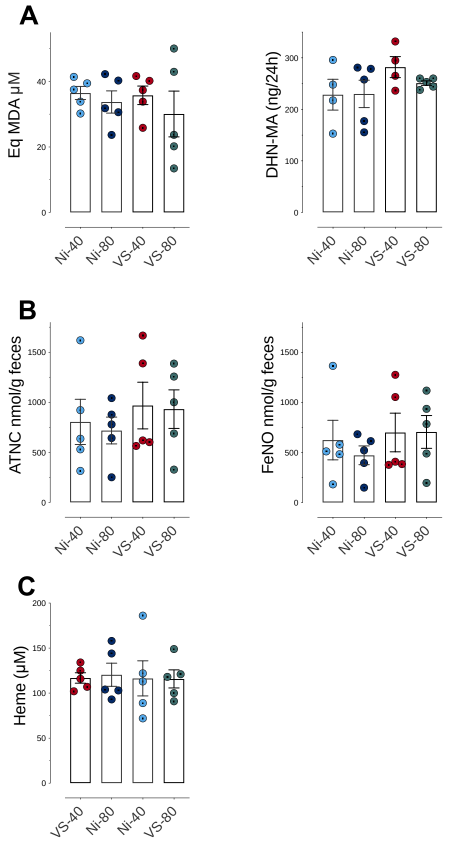


**Figure S1. Impact of sodium nitrites levels and vegetable stocks in cooked ham models on fecal and urinary biomarkers of lipid peroxidation and NOCs formation**

**A**-Lipid peroxidation measured as TBARS (MDA equivalents, mM) in fecal water and DHN-MA in urine of 24h (ng/vol of 24h). **B**-Nitroso-compounds in fecal water measured as total NOCs (ATNC), as nitrosyl iron (FeNO), (nmol/gd of feces) **C**-Heme in fecal water (µM) Data were represented using scatter plots with bar (mean ± sem), * $p \leq 0.05$; ** $p \leq 0.01$.

#### 2. ***Biochemical characteristics of cooked ham models used for the CRC animal study***

##### *2.1 Supplementary experimental section*

###### *2.1.1 Iron, zinc, heme iron and nitrosylheme.*

Total heme iron content was assessed in the form of acid hematin and was extracted in acidic acetone[3]. The samples were mixed in the dark for 2 hours and filtered through 0.45 µm regenerated cellulose membranes (Interchim®, France). Then, the absorbance was measured at 512 nm with an absorption coefficient of 9.52 $mM^{-1}cm^{-1}$ with a Jasco V770 spectrophotometer[4] (Jasco, Oklahoma City, USA). Nitrosylheme content was assessed by diluting the samples in an acetone/water mixture. Samples were then mixed in the dark for 15 minutes and filtered through 0.45 µm filters (Interchim®, France). Nitrosylheme absorbance was measured at 540 nm using an absorption coefficient of 11.3 $mM^{-1}cm^{-1}$ [4]. Nitrosylation was expressed as the percentage of nitrosylated iron to total heme iron. The concentration of total non-heme iron, $Fe^{2+}$, and $Fe^{3+}$ was determined using the ferrozine method, according to Stolze, Dadak, Liu and Nohl[5]. Briefly, samples were diluted three times in citrate buffer (15 mM citrate, 140 mM NaCl, pH 7.4) and centrifuged with Vivapsin® systems (5 kDa cut-off) for 75 minutes at 18°C and 4,000 rpm to remove heminic iron. Then, the absorbance was measured at 562 nm on a multiskan spectrum from Thermo Scientific (Waltham, USA). A standard curve was constructed using ferrous sulfate ($FeSO_4$) containing ascorbic acid. The results are expressed in mg/kg. In addition, the trace elements Fe, and Zn were quantified by inductively coupled plasma spectrometry (ICP-AES) following the Poitevin method[6]. The results are expressed in mg/kg.

***2.1.2 Lipid peroxidation.***

One g of samples was mixed in 9 mL buffer (0.15 M KCl, 0.1 mM BHT) using a Polytron PT 2100 (Kinematica AG, Switzerland). The samples were centrifuged at 13,000 g for 10 minutes, and the supernatant was collected. A solution of thiobarbituric acid (TBA) prepared in glacial acetic acid and water was added to the samples to reach a 4-fold dilution. The reaction was started by incubating the samples at 95°C for 60 minutes, after which they were cooled on ice for 10 minutes. The malondialdehyde (MDA)-TBA adducts were measured at 532 nm and 760 nm (for turbidity) with a spectrophotometer (Jasco V770, Oklahoma City, USA). A standard curve was constructed using 1,1,3,3-tetraethoxypropane (TEP) and results are expressed in µg MDA/g.

***2.1.3. Residual nitrite, residual nitrate, nitrosothiols, non-volatile and volatile nitrosamines.***

Nitroso-compounds were quantified as described previously[3]. Briefly, 2 g of cooked ham model were homogenised in 10 mL water containing 2.4 mL of 2% NaOH using a Polytron PT 2100 (Kinematica AG, Switzerland). The samples were then further diluted in water, and the pH was adjusted to 8 with HCl. Then, the samples were heated at 50°C for 15 minutes in a water bath. The heat extraction was followed by deproteinization and ultrafiltration of the samples achieved with Vivapsin® systems (5 kDa cut-off). Samples were centrifuged at 4,000 rpm, 16°C for 75 minutes (SL 40R centrifuge; Thermo-Scientific, USA). The supernatants were eliminated and the filtrates were used for following assays.

Nitrite and nitrate ion contents were then determined in the filtrates using the Griess reaction with a Sigma-Aldrich colorimetric assay kit (23479-1KT-F). Absorbance was measured on a microplate at 540 nm on a MULTISKAN SPECTRUM spectrophotometer from Thermo Scientific (Waltham, USA). Residual nitrite and nitrate were expressed in mg/kg.

The filtrates were added to 1% (*w*/*v*) $HgCl_2$ to break the *S*-NO bond. Samples were incubated for 10 minutes in the dark, after which excess $HgCl_2$ was removed by filtration through 0.22 µm filters (Sartorius, Germany). Nitrite ion content release was measured above using the

Griess reaction with the Sigma-Aldrich colorimetric assay kit, and the difference between this measurement and the level of nitrite ions initially present in the sample (Section 1.1.3.1) gave the nitrosothiol content (expressed in mg/kg).

The *N*-NO bond of non-volatile nitrosamines was broken during a 2-hour UV treatment (LF 215.S, 254 nm, 2x15 W). Aliquots were taken after 1, 15, 30, 60, and 120 minutes to construct a kinetic curve of combined nitrosothiols and nitrosamines content. The contents of both nitrite and nitrate was measured using the Griess reaction. The concentration of [$NO_2^-$ + $NO_3^-$] was plotted versus the inverse of time and extrapolation to 1/t = 0, obtained by fitting a second-degree polynomial function, giving the maximum rate of combined nitrite and nitrate released after the UV treatment. The subtraction of nitrosothiol content, obtained as described above, gave the non-volatile *N*-nitrosamine and the nitrosamide contents (Apparent Total *N*-Nitroso Compounds, ATNC). The results were expressed in mg/kg.

Regarding volatile nitrosamines, assays were carried out and quantified by the Eurofins company (https://www.eurofins.fr/). Five nitrosamines (*N*-Nitrosodimethylamine (NDMA), *N*-Nitrosomethylethylamine (NMEA), *N*-Nitrosodiethylamine (NDEA), *N*-nitrosodiisobutylamine (NDiBA), *N*-Nitrosodibutylamine (NDBA)) were quantified in triplicate on cooked ham models stored at -80°C until analysis. Quantification was performed by LC-(APCI)MS/MS.

##### *2.2 Supplementary result section*

| | **Iron** | **Zinc** |
|---|---|---|
| **PRE** | 1.15 ± 0.02 | 2.92 ± 0.20 |
| **Ni-120** | 1.25± 0.14 | 2.93 ± 0.19 |

**Table S1. Quantification of total iron and zinc by ICP AES in PRE and Ni-120 cooked ham models, expressed in mg/kg**

Data were mean ± SD

The results presented in Table S1 show an equivalent zinc concentration between the two cooked ham models.

| | n | NDMA | NMEA | NDEA | NDiBA | NDBA | NDMA + NDEA |
|---|---|---|---|---|---|---|---|
| Ni-120 | 3 | <1.0 | <1.0 | <1.0 | <1.0 | <1.0 | <1 |
| VS | 3 | <1.0 | <1.0 | <1.0 | <1.0 | <1.0 | <1 |
| PRE | 3 | <1.0 | <1.0 | <1.0 | <1.0 | <1.0 | <1 |
| YE | 3 | <1.0 | <1.0 | <1.0 | <1.0 | <1.0 | <1 |

**Table S2. Quantification of five volatile *N*-nitrosamines (µg/kg) in Ni-120 and the three alternatives (VS, PRE and YE).** *N*-Nitrosodimethylamine (NDMA), *N*-Nitrosomethylethylamine (NMEA), *N*-Nitrosodiethylamine (NDEA), *N*-nitrosodiisobutylamine (NDiBA), *N*-Nitrosodibutylamine (NDBA)

The results presented in Table S2 show results below the quantification limit of the method.

#### 3. *Listeria monocytogenes growth assay in sliced cooked ham model products*

##### *3.1 Supplementary experimental section*

To follow the growth of *L. monocytogenes*, a sliced cooked ham model, specifically produced for this assay, was used: 6 different recipes with various amounts of added nitrite (Ni-120, Ni-90 and Ni-0) or alternatives to nitrite (VS, PRE and YE) were tested (Table S3).

###### *3.1.1 Specific production of the sliced cooked ham model products*

These products were composed of 2.7 kg ground pork supplemented with a specific brine solution as described below. The sliced cooked ham model products were made in a pilot-scale unit (IFIP, France). They were prepared with fresh boned and derinded pork shoulders. Further trimming and cutting were performed, and the pieces of meat were ground through a 20 mm plate of a grinder (DRC 98, PSV, France). In each experiment, the ground pork was divided into as many recipes as necessary, then mixed with appropriate concentrations of sodium nitrite (0, 90 or 120 mg/kg) or alternatives, NaCl (18 or 20 g/kg), sodium ascorbate (0 or 300 mg/kg), dextrose (0 or 5 g/kg) and/or saccharose (0 or 1 g/kg) using a vacuum mixer (CDH, France) at a constant speed of 350 rpm for 10 min. The different starters used in the VS and YE recipes were stored at -20°C and prepared according to the manufacturer's instructions. In all recipes, water was added in order to have a 10% brine rate (Table S3).

For each recipe, two 2.7 kg cooked ham model products were manufactured. To this end, the cured ground pork was dispensed into two plastic molds and vacuum packed in heat shrink bags (43 µm-multilayer bags including polyamide, polyethylene and ethylene vinyl alcohol; $O_2$ transmission rate, 12 $cm^3/m^2$ in 24 h at 20°C and 90% relative humidity; moisture vapor transmission rate, 15 $g/m^2$ in 24 h at 38°C and 90% relative humidity) (SURVID VAC,

Soussana, France) using a Multivac A300 packaging machine (Multivac, France). They were then stored overnight at 3°C to simulate vacuum tumbling before thermal processing.

##### *3.1.2* Thermal processing, slicing and storage conditions of the cooked ham model products

Vacuum packed products were then subjected to thermal treatment classically used in the meat processing industry. Cooking and cooling processes were performed in a saturated steam oven (SelfCookingCenter® 61, Rational, Belgium). Temperature of the cooking and cooling processes was monitored using two PT100 temperature probes, one inserted into a 2.7 kg product and another one placed in ambience of the oven chamber (Fig S2). All values were registered by using a datalogger Almemo® 2590 (Ahlborn, Germany). The core temperature of the processed meat product reached 67°C after 514 min followed by a cooling process that required 851 min from 67°C to 4°C (representative of a pasteurization value $PV_{70,10}$ of 60 min). It is important to note that the thermal processing allowed the development of both the YE and VS starters.

Once cooled at 4°C, cooked ham products were unpacked then sliced using a TIP350 slicer (Simplex, France). Sliced cooked ham samples were then surface inoculated with the cocktail of three *L. monocytogenes* strains as described in the § 3.1.3, then packed under protective atmosphere (gas mixture including 50 % $CO_2$ and 50 % $N_2$) in polyamide (20 µm) and polyethylene (70 µm) bags ($O_2$ transmission rate, 50 $cm^3/m^2$ in 24 h at 23°C and 85% relative humidity; moisture vapor transmission rate, 2.60 g/m2 in 24 h at 23°C and 85% relative humidity) (Soussana, France) by using the Multivac A300 packaging machine. These samples were stored under reasonably foreseeable conditions (time and temperature) that could be encountered along the supply chain i.e., 14 days at 4°C (cold storage chamber) and 35 days at 8°C (distributor shelving and domestic storage in refrigerator).

##### *3.1.3 Listeria monocytogenes* strains and inoculation of the sliced cooked ham model samples

A cocktail of 3 *Listeria monocytogenes* i.e., Lm176 (isolated from cooked ham, clonal complex 2), Lm212 (isolated from cooked ham, clonal complex 101) and Lm004 (isolated from “rillettes” involved in a previous outbreak of listeriosis, clonal complex 1) (IFIP collection) was used to inoculate the sliced cooked ham model samples from the different recipes. The use of several strains mixed in a cocktail is recommended by the ISO 20976-1 standard for estimation of growth potential so that variations among strains are considered.

These strains were stored at -80°C on cryobeads. For each strain, a preculture was prepared in BHI broth (bioMérieux, France) kept at 37°C overnight. Precultures were diluted 100-fold in BHI broth and subsequently incubated at 8°C for 5 days. After incubation, culture of each *L. monocytogenes* strain was enumerated on both non-selective Tryptone Soya Yeast Extract Agar (TSAYE) (Oxoid, France) and selective chromogenic AL agar (Agar Listeria according to Ottaviani and Agosti) (Bio-Rad, France) plates following an overnight incubation at 37°C. A cocktail of the three *L. monocytogenes* strains in equal proportions was then prepared in saline solution, which was used to inoculate the different sliced cooked ham model samples (i.e. 2 cooked ham slices of about 30 g each per sample) at 2 $\log_{10}$ CFU/g (mean inoculum value ± standard deviation (SD) of 2.0 ± 0.12 $\log_{10}$ CFU/g, all 3 experiments included).

##### *3.1.4 Microbiological analysis of the sliced cooked ham model samples*

At each sampling date (D0, D07, D14, D17, D22, D28, D35 and D49), three samples per recipe were analyzed for the enumeration of *L. monocytogenes* using the BRD 07/17 - 01/09 standard method. Briefly, each 60 g sample was four-fold diluted in buffered peptone water (Bio-Rad, France) and stomacher at medium speed for 1 min. One hundred microliters of this suspension and/or of the appropriate tenfold serial dilutions were plated on AL agar plates which were incubated at 30°C for 48 h before counting colonies. The quantification limit of this analysis was 4 CFU/g. Moreover, one uninoculated sample per recipe was analyzed for natural contamination of *L. monocytogenes* using the BRD 07/16 - 01/09 standard method for *L. monocytogenes* detection at the beginning and at the end of the storage period. Briefly, samples were enriched for 24 h at 30°C in half Fraser broth followed by subculture on AL agar plates which were incubated at 37°C for 24 h. In case of suspect colonies, confirmation test was performed using selective chromogenic RAPID'*Listeria* spp. agar plates (Bio-Rad, France) which were incubated for 24 h at 37°C. Raw material samples tested in the present study were all negative for *L. monocytogenes* (data not shown).

For each recipe, lactic acid bacteria were enumerated in triplicate (from 3 different samples/recipe) at D0 and D49 by plating appropriate dilutions on MRS medium (bioMérieux, France) after a 48 h-incubation at 30 °C. The quantification limit of this analysis was 10 CFU/g. Results are shown in the Table S5.

##### *3.1.5 Physico-chemical analysis of the sliced cooked ham model samples*

For each recipe, pH and $a_w$ values were determined according to the NF ISO 21807:2005 and NF V04–408 standards, respectively. Aw and pH measurements were performed on three sliced cooked ham samples per recipe at D0, D22 and D49 using an AQUALAB 4TEV (Meter,

Germany) and a FiveGo (Mettler-Toledo, Switzerland) apparatus connected to a puncture electrode LE427-IP67 (Mettler-Toledo, Switzerland), respectively. Oxygen and carbon dioxide were measured using an $O_2$ and $CO_2$ head space gas analyzer, Checkmate 3 (Dansensor, France) at D0 and D49. At the same sampling dates, residual nitrite and nitrate contents were measured in 3 samples per recipe by using a flow injection analysis. The sodium contents were determined on 3 samples per recipe at D0 following dry mineralization and atomic absorption spectroscopy. Nitrite, nitrate and sodium contents analyses were performed at Actalia laboratory (Villers Bocage, France). Colour measurements were carried out with a CM 600 spectrophotometer (Minolta, Japan) configurated as follows: measurement area of 8 mm in diameter, illuminant D65, geometry d/8°. The value considered was the red hue angle (H*). The measurements were carried out in triplicate on the surface of 3 sliced cooked ham samples per recipe at D0 (after slicing) and D49. Lipid oxidation was measured by the thiobarbituric acid reactive substances (TBARS) content using the method of Mercier et al.[7], with slight modifications. The product (700 mg) was dispensed in plastic tubs containing ceramic beads (CK28) and homogenized with 7 mL buffer (0,15 M KCl, 0,1 mM BHT). Before and during the tissue homogenization, the temperature was maintained at 4°C due to a device based on dry ice sublimation. Extract obtained (1 mL) was incubated with 1% (w/v) 2-thiobarbituric acid in 50 mM NaOH (0.50 mL) and 2.8% (w/v) trichloroacetic acid (0.50 mL) in a boiling water bath for 10 min. After cooling to room temperature (30 min), the pink chromogen (MDA-TBA adducts) was extracted into n-butanol (2.0 mL) and its absorbance measured at 535 nm and 760 nm (for turbidity) with a spectrophotometer (Uvikon). A standard curve was prepared using a 1,1,3,3-tetraethoxypropane (TEP) and results were expressed as mg MDA equivalent (MDA eq)/ kg sample. For each experimental assay and recipe, measurements were performed in triplicate on a pool of 3 cooked ham model samples from D49. Results are displayed in the Table S4.

##### *3.1.6 Statistical analysis*

For each independent experiment, growth potential $(\delta)$ was calculated according to the equation $(\delta) = log_{10\,t} - \overline{log_{10\,\iota}}$ (adapted from the ISO 20976-1:2019 standard), where $log_{10\,t}$ is the value obtained for one sample at a given sampling date and $\overline{log_{10}\,\iota}$ is the mean value obtained for the three replicate samples analyzed at D0. When δ was higher than 0.5 $log_{10}$ CFU/g the sliced cooked ham model sample was considered permissive to the growth of *L. monocytogenes*.

The effect of the recipe on growth potentials of *L. monocytogenes* on sliced cooked ham model samples was assessed by using the ANOVA test followed, when significant, by pairwise comparison using the estimated marginal means followed by Bonferroni correction. All statistical analyses were performed using the RStudio statistical software (version 2021.09.0 Build 351—2009–2021).

| Ingredients or additives (g/kg of cured meat) | Recipes | | | | | |
|---|---|---|---|---|---|---|
| | **Ni-120** | **Ni-90** | **Ni-0** | **VS** | **PRE** | **YE** |
| Sodium nitrite ($NaNO_2$) | 0.12 | 0.09 | 0 | 0 | 0 | 0 |
| Nitrated broth NAT 223 | 0 | 0 | 0 | 4 | 0 | 0 |
| Starters CS300 | 0 | 0 | 0 | 0.25 | 0 | 0 |
| Polyphenol-rich and ascorbic acid rich extract | 0 | 0 | 0 | 0 | 10 | 0 |
| Yeast extract | 0 | 0 | 0 | 0 | 0 | 5 |
| Starters | 0 | 0 | 0 | 0 | 0 | 0.2 |
| Total sodium chloride | 18 | 18 | 18 | 20 | 18 | 20 |
| Sodium ascorbate | 0.3 | 0.3 | 0.3 | 0.3 | 0 | 0 |
| Dextrose | 5 | 5 | 5 | 5 | 5 | 0 |
| Saccharose | 0 | 0 | 0 | 0 | 0 | 1 |
| Water (mL/kg of meat) | 75 | 74 | 74 | 67 | 64 | 68 |

**Table S3: List of the ingredients and additives** included in the 6 recipes tested to produced cooked ham models. See main manuscript "experimental section" for component origin.

##### *3.2 Supplementary Result Section*

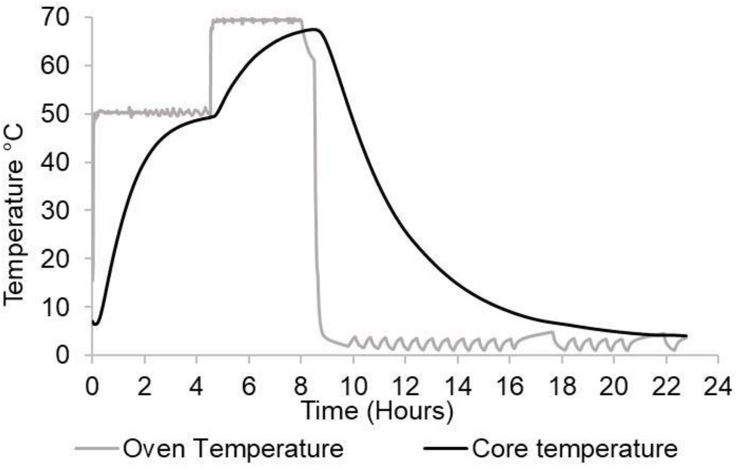


**Figure S2: Heat treatment kinetics** obtained in ambience of the oven chamber (grey) and into the cooked ham model product (black) for the first experimental trial as an example

| Recipe | pH | | $a_w$ | | NaCl (%) | | $NaNO_2$ (mg/kg) | | $NaNO_3$ (mg/kg) | | TBARS (mg MDA eq /kg) | | Red tint angle ° | |
|---|---|---|---|---|---|---|---|---|---|---|---|---|---|---|
| | D0 | D49 | D0 | D49 | D0 | D49 | D0 | D49 | D0 | D49 | D0 | D49 | D0 | D49 |
| **Ni-120** | 6.12 ± 0.047 | 5.96 ± 0.230 | 0.979 ± 0.004 | 0.980 ± 0.008 | 1,78 ± 0,05 | *ND* | 34.8 ± 7.1 | 8.4 ± 0.15 | 18.5 ± 2.44 | 33.3 ± 2.45 | *ND* | 0.312 ± 0.40 | 48.4 ± 3.03 | 48.4 ± 3.94 |
| **Ni-90** | 6.12 ± 0.043 | 5.88 ± 0.227 | 0.978 ± 0.003 | 0.979 ± 0.009 | 1,77 ± 0,06 | *ND* | 19.2 ± 6.11 | 3.2 ± 1.85 | 12.8 ± 1.60 | 23.1 ± 5.28 | *ND* | 0.321 ±0.035 | 48.4 ± 3.52 | 50.5 ± 2.86 |
| **Ni-0** | 6.11 ± 0.069 | 5.92 ± 0.231 | 0.98 ± 0.003 | 0.978 ± 0.006 | 1,8 ± 0,05 | *ND* | <1.5 | <1.5 | <6.9 | <6.9 | *ND* | 1.613 ±0.160 | 19.0 ± 2.36 | 12.3 ± 3.46 |
| **VS** | 6.11 ± 0.073 | 5.86 ± 0.219 | 0.975 ± 0.003 | 0.982 ± 0.005 | 1,91 ± 0,06 | *ND* | 15.4 ± 5.88 | <1.5 | 10.6 ± 5.58 | 17.7 ± 0.74 | *ND* | 0.300 ± 0.035 | 45.6 ± 3.93 | 49.1 ± 2.92 |
| **PRE** | 6.07 ± 0.090 | 5.84 ± 0.087 | 0.975 ± 0.005 | 0.977 ± 0.006 | 2,04 ± 0,07 | *ND* | <1.5 | <1.5 | 7.4 ± 0.38 | <6.9 | *ND* | 0.316 ±0.045 | 48.6 ± 3.43 | 48.0 ± 4.6 |
| **YE** | 6.10 ± 0.046 | 5.99 ± 0.300 | 0.976 ± 0.002 | 0.978 ± 0.004 | 2,07 ± 0,08 | *ND* | <1.5 | <1.5 | <6.9 | <6.9 | *ND* | 2.977 ±1.320 | 27.5 ± 2.54 | 16.0 ± 4.26 |

**Table S4**: **Physicochemical properties of the cooked ham model products** from the different recipes (mean values ± SD obtained from all 3 independent experiments).

| Recipe | Lactic acid bacteria ($Log_{10}$ CFU/g) | |
|---|---|---|
| | D0 | D49 |
| **Ni-120** | 0.0 ± 0.00 | 7.8 ± 0.67 |
| **VS** | 0.4 ± 0.65 | 7.8 ± 0.87 |
| **PRE** | 0.6 ± 1.00 | 8.2 ± 0.38 |
| **YE** | 0.0 ± 0.00 | 7.9 ± 0.98 |
| **Ni-90** | 0.3 ± 0.50 | 8.2 ± 1.16 |
| **Ni-0** | 0.4 ± 0.61 | 7.1 ± 1.14 |

**Table S5**: **Lactic acid bacteria populations** ($Log_{10}$ CFU/g) enumerated in the cooked ham model products from the different recipes (mean values ± SD obtained from all 3 independent experiments).

Cooked ham model samples used in the microbiological assays exhibited typical pH (mean values from 6.07 to 6.12 regardless of the recipes) and $a_w$ (mean values from 0.975 to 0.980 regardless of the recipes) values at D0. The $a_w$ of the products remained steady during the 49-day storage whereas the pH slightly decreased (minimal mean value of 5.84 units regardless of the recipe). The proliferation of lactic acid bacteria (Table S4) can be associated to the decrease in the measured pH levels. All pH and $a_w$ values measured during the present study were permissive to the growth of *L. monocytogenes* (ANSES, 2021). The nitrate salt, nitrite salt and NaCl levels measured were consistent with the additive doses in corresponding recipes (Table S3 and S5).

#### *4. Fecal N-nitroso compounds (NOC) assay*

##### *4.1 Supplementary Experimental Section*

###### *4.1.1 Quantification of fecal NOC*

Fecal water samples were thawed on ice, followed by centrifugation at 16.000 g and 4°C for 20 min. Afterwards, the supernatant was transferred into a fresh tube and diluted with ultrapure water if necessary. Each fecal water sample was measured in triplicate after three individual pretreatments incubated on ice: (I) 100 µL of diluted sample was mixed with 200 µL sulfanilamide (30 mg/mL in 1 M HCl), incubated for 5 min and further diluted with 100 µL ultrapure water; (II) 100 µL of diluted sample was mixed with 100 µL $HgCl_2$ (14.5 mg/mL in ultrapure water) and incubated for 30 min, followed by the addition of 200 µL sulfanilamide and an incubation for 5 min; (III) 100 µL of diluted sample was mixed with 100 µL $K_3Fe(CN)_6$ (38 mg/mL in ultrapure water) and incubated for 30 min, followed by the addition of 200 µL sulfanilamide and an incubation for 5 min. After pretreatment, 100 µL of each sample was injected into the purge vessel, containing 10.5 mL of the following reaction solution: 2 mL KI (50 mg/mL in ultrapure water), 400 µL $CuSO_4$ (50 mg/mL in ultrapure water), 8 mL acetic acid and 100 µL antifoam. The reaction solution was kept at 60°C and changed regularly after measuring five pretreated samples in triplicate. The area under curve was quantified by sodium nitrite standards (10 - 250 nM) dissolved in ultrapure water. For differentiation of apparent total *N*-nitroso compounds (ATNC) in *S*-nitrosothiols (RSNO), nitrosyl iron (FeNO) and residual *N*-nitroso compounds (RNNO), the contents were calculated individually by the results of pretreatments. Pretreatment I allowed calculation of ATNC content. RSNO contents were obtained by subtraction of the results of pretreatment II from the results of pretreatment I. FeNO contents were calculated similarly by subtraction of the results of pretreatment III from the results of pretreatment I. Finally, RNNO contents were determined by the substraction of the sum of RSNO and FeNO from ATNC. All results are shown as nanomole per gram feces.

###### *4.1.2 Test of interference with polyphenols in fecal water*

To check whether polyphenols and ascorbic acid present in the PRE formulation may interfere with the NOC assay and generate false-positive results, pooled fecal waters (100µL) from Ni-0 ham fed rats were added with 0.2 mg of epigallocatechine gallate (EGCG), 0.8 mg ferulic acid and 0.5 mg ascorbic acid (Sigma-Aldrich) and analysed for Total NOCs with the method described in 4.1.1. The mixture is called “FEA”

##### *4.2 Supplementary Result Section*

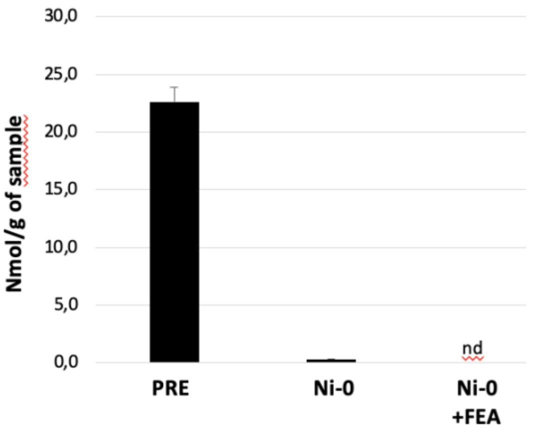


**Figure S3: ATNC concentration in fecal water** from PRE ham fed rats, from Ni-0 ham fed rats and fecal water of from Ni-0 ham fed rats supplemented in epigallocatechine gallate, ferulic acid and ascorbic acid (Ni-0 + FEA). Mean values ± SD

This assay, independent of the analyses presented in figure 6 of the main manuscript, confirmed a high presence of ATNC in the PRE fecal water and their absence in the Ni-0 fecal water. We also demonstrated that the overloading of Ni-0 fecal water with polyphenols and ascorbic acid present in the PRE formulation did not interfere with the assay and did not generate false positive results with our NOC analysis method.

#### *5. Cooked ham model N-nitroso compounds (NOC) assay*

##### *5.1 Supplementary Experimental Section*

Nitrosyl heme and total heme were determined in Ni-0 ham (Ni-0) and the same ham loaded with different concentrations of epigallocatechine gallate (EGCG) (5 mg/mL, 10 mg/mL, 20 mg/mL). 1g of Ni-0 ham was grounded with 1 mL of water (Ni-0) and 1g of the same ham was grounded in 1mL solution containing 100 µL EGCG 5 mg/mL in $H_2O$ or 100 µL EGCG 10 mg/mL in $H_2O$ or 100 µl EGCG 20 mg/mL in $H_2O$ +650 µL $H_2O$ (EGCG5, EGCG10, EGCG20).

1g of Ni-0 ham was grounded with 1 mL of water (Ni-0) and 1g of Ni-0 ham was grounded in 1 mL solution containing 100 µL EGCG 20 mg/mL in $H_2O$; 150 µL ferulic acid 53 mg/mL in ethanol; 100 µL ascorbic acid 50 mg/mL in water; 650 µL $H_2O$ (Ni-0 + FEA).

##### *5.2 Supplementary Result Section*

| | Total heme iron mM | Fe NO mM | % Fe-NO |
|---|---|---|---|
| Ni-0 | 0.14 ± 0.013 | 0.0012 ± 0.000 | 0.5 |
| ECGC 5 | 0.11 ± 0.01 | 0.0014 ± 0.000 | 1.3 |
| ECGC 10 | 0.11 ± 0.007 | 0.0023 ± 0.0006 | 2.2 |
| ECGC 20 | 0.12 ± 0.004 | 0.0049 ± 0.0002 | 4.1 |
| $p < 0.05$ | NS | * | * |

**Table S6**: **Total heme iron and nitrosyl heme iron (FeNO)** in Ni-0 ham (Ni-0) and the same ham loaded with different concentrations of epigallocatechine gallate (EGCG) (5 mg/mL, 10 mg/mL, 20 mg/mL).

This assay showed interference in the determination of nitrosyl heme by colorimetric method (Hornsey) when EGCG is added, while it was not the case for total heme iron.

| | Total heme iron mM | FeNO mM | % Fe- NO |
|---|---|---|---|
| Ni-0 | 0.13 ± 0.02 | 0.0018 ± 0.0004 | 1.1 |
| Ni-0 + FEA | 0.21 ± 0.01 | 0.0217 ± 0.00005 | 10.6 |

**Table S7**: **Total heme iron and nitrosyl heme iron (FeNO)** in Ni-0 ham (Ni-0) and the same ham loaded with a mix of epigallocatechine gallate, ferulic acid and ascorbic acid (Ni-0 + FEA). (mean value ± SD)

This assay showed interference in the determination of total heme iron and nitrosyl heme by colorimetric method (Hornsey) when a mix of epigallocatechine gallate, ferulic acid and ascorbic acid was added. The total heme iron is over estimated by 60% while for nitrosyl iron the overestimation is multiply by 8. Lastly the % of nitrosyl heme is approximately overestimated by 10.

| | $NO_2$ mg/kg | $NO_3$ mg/kg | RSNO mg/kg | RNNO mg/kg |
|---|---|---|---|---|
| Ni-0 | 0.05 ± 0.04 | 3.76 ± 0.09 | 0.23 ± 0.08 | 0.06 ± 0.04 |
| Ni-0 + FEA | 0 ± 0 | 1.97 ± 0.24 | 0.05 ± 0.04 | 0.33 ± 0.10 |

**Table S8**: **Colorimetric determination of $NO_2$, $NO_3$, RSNO and RNNO** according to Bonifacie et al 2021[3] using the Griess reagent, in Ni-0 ham (Ni-0) and the same ham loaded with a mix of epigallocatechine gallate (EGCG), ferulic acid and ascorbic acid (Ni-0 +FEA). 1g of Ni-0 ham was grounded with 1 mL of water (Ni-0) and 1g of Ni-0 ham was grounded in 1mL solution containing 100 µL d'EGCG 20 mg/mL in $H_2O$; 150µL ferulic acid 53 mg/mL in ethanol; 100 µL ascorbic acid 50 mg/mL in water; 650 µL $H_2O$ (Ni-0 + FEA). Then nitroso-compounds were quantified as described previously[3] in section 2.1.3.

This assay did not show any interference in the determination of $NO_2$, NO, RSNO, RNNO by the colorimetric method of Griess when a mix of epigallocatechine gallate, ferulic acid and ascorbic acid was added. The level of RSNO and RNNO even significantly different were under the threshold sensitivity of 1 mg/kg.

#### 5. *Colon mucosa gene expression assays on samples of the 100days-study with different concentrations of sodium nitrite (0, 90 and 120 mg/kg) or sodium nitrite alternatives in cooked ham models.*

##### *5.1 Supplementary Experimental Section*

*Gene expression assays in colon mucosa: Total* cellular RNA was extracted with Tri reagent (Molecular Research Center). Total RNA samples (1 µg) were then reverse-transcribed with the iScript™ Reverse Transcription Supermix (Bio-Rad) for real-time quantitative polymerase chain reaction (qPCR) analyses. The primers for Sybr Green assays are presented in Table S5. Amplifications were performed on a ViiA 7 Real-Time PCR System (Applied Biosystems). The qPCR data were normalized to the level of the RNA Polymerase II Subunit A (POLR2A) messenger RNA (mRNA) and analyzed by the LinRegPCR v.11 software.

| Gene | Alias | sequence primer Forward | sequence primer Reverse |
|---|---|---|---|
| **Polr2a** | RNA Polymerase II Subunit A | CGGCGTCCTGAGTCCG | AACTTGGGGGACTAATGGATCC |
| **Aldh2** | Aldehyde Dehydrogenase 2 Family Member | GCCGCAGACCGTGGTTACT | CGATGGTCATGCCATCTTTG |
| **Aldh3a2** | Aldehyde Dehydrogenase 3 Family Member A2 | TCTGAGGCAGCGGTTTGATC | CCAACTGCGGTGTTTCCTGT |
| **Akr1b8** | Aldo-Keto Reductase Family 1 Member B8 | GCCGCCAAGCACAAGAAAA | CCTCTGGATGTGGAACCGAAT |
| **Akr1b10** | Aldo-Keto Reductase Family 1 Member B10 | CCTGGGCACCTGGAAGACT | CATCAATGGCTGCCTTCACA |
| **Nqo1** | NAD(P)H Quinone Dehydrogenase 1 | CGCAGAGAGGACATCATTCA | GTGGTGATGGAAAGCAAGGT |
| **Keap1** | Kelch Like ECH Associated Protein 1 | CTGCATCCACCACAGCAGCGT | GTGCAGCACACAGACCCCGGC |
| **Hmox1** | Heme Oxygenase 1 | CAACCCCACCAAGTTCAAACA | AGGCGGTCTTAGCCTCTTCTG |
| **Gclc** | Glutamate-Cysteine Ligase Catalytic Subunit | GTCTTCAGGTGACATTCCAAGC | TGTTCTTCAGGGGCTCCAGTC |
| **Gsta4** | Glutathione S-Transferase Alpha 4 | GAAGTTCTAGTGACAGCGTGCTTTA | TGTAGCTGCTGCTGTGATTGG |
| **Slc7a11** | Solute Carrier Family 7 Member 11 (xCT) | CCTGGCATTTGGACGCTACAT | TCAGAATTGCTGTGAGCTTGCA |
| **Txn** | Thioredoxin (TRX1) | CTTTCATTCCCTCTGTGACAAGTATTC | TTATAGAACTGGAAGGTCGGCATG |
| **Cat** | Catalase | ACGAGATGGCACACTTTGACAG | TGGGTTTCTCTTCTGGCTATGG |
| **Sod1** | Superoxide Dismutase 1 | GCGGTCCAGCGGATGA | GTCCTTTCCAGCAGCCACAT |
| **Ptgs2** | Prostaglandin-Endoperoxide Synthase 2 (Cox2) | AGATCAGAAGCGAGGACCT | CCATCCTGGAAAAGTCGAA |
| **Cbr1** | Carbonyl Reductase 1 | CAAGGAGCTACTCCCTATAATAA | CATGCCGCTTGATACATTC |
| **Ptgr1** | Prostaglandin Reductase 1 | TACCACTGTCTTTGGCTTTGGG | CCCTTCAAGCCACAGATGTCAA |
| **Gss** | Glutathione Synthetase | GCATCTCGGAGCTGGGTATTT | TCACAAGTGTTGTTCCCTGTCT |
| **Gsr** | Glutathione-Disulfide Reductase | GAAGATGGTTTGTGCCAACAAAGAG | GCATATGGATGCCAACCACCTT |

**Table S9: Primer sequences for RT-qPCR analysis.**

#### *5.2 Supplementary results*

Reduction or removal of sodium nitrite concentration had low or no impact on colon mucosal detoxification activities. The most striking effect, although not significant, is the dose-dependent increase in the expression of Nqo1, and trends to a decrease for Gsta4, Akr1b10, Cbr1, Sod1 in the case of sodium nitrite removal (Figure S4).

The effect of the alternatives appeared to be weaker with only one notable and significant effect of YE cooked ham model that induced a significant decrease in the expression of Hmox1 compared to cooked ham models VS and PRE and a significant increase in the expression of Cox2 compared to the reference cooked ham model Ni-120 and VS (Figure S5).

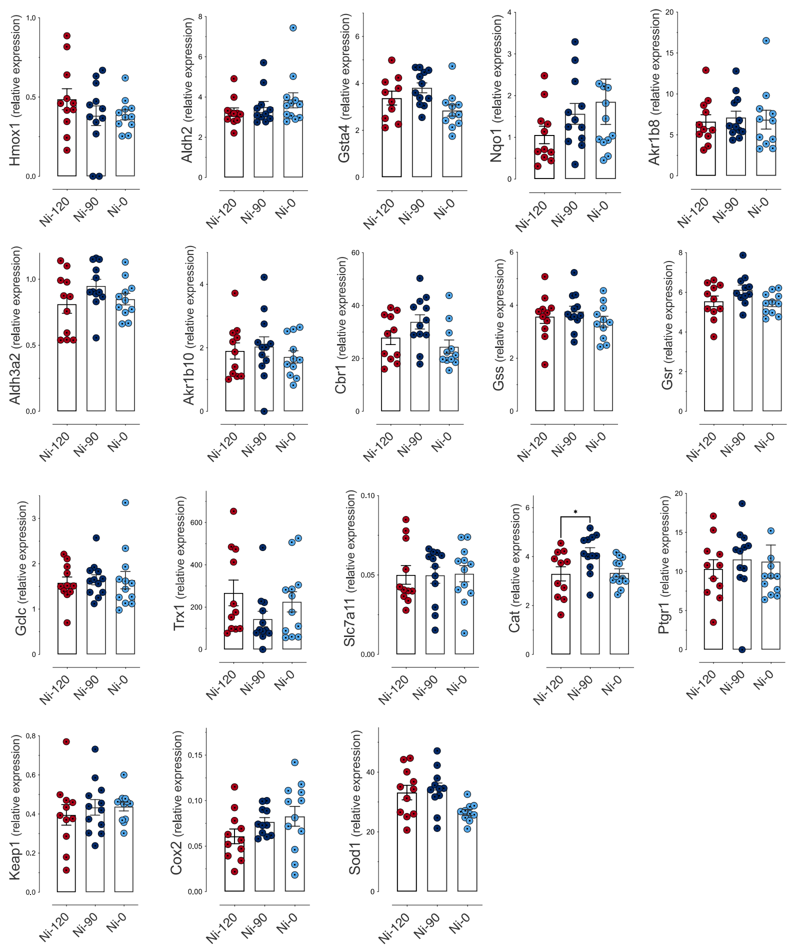


**Figure S4. Impact of sodium nitrite concentrations in cooked ham models (0 *vs* 90 *vs* 120 mg/kg) on gene expression in rat colon mucosa.** Data were represented using scatter plots with bar (mean ± sem), * $p \leq 0.05$. Genes full names are shown in sup. Table S9.

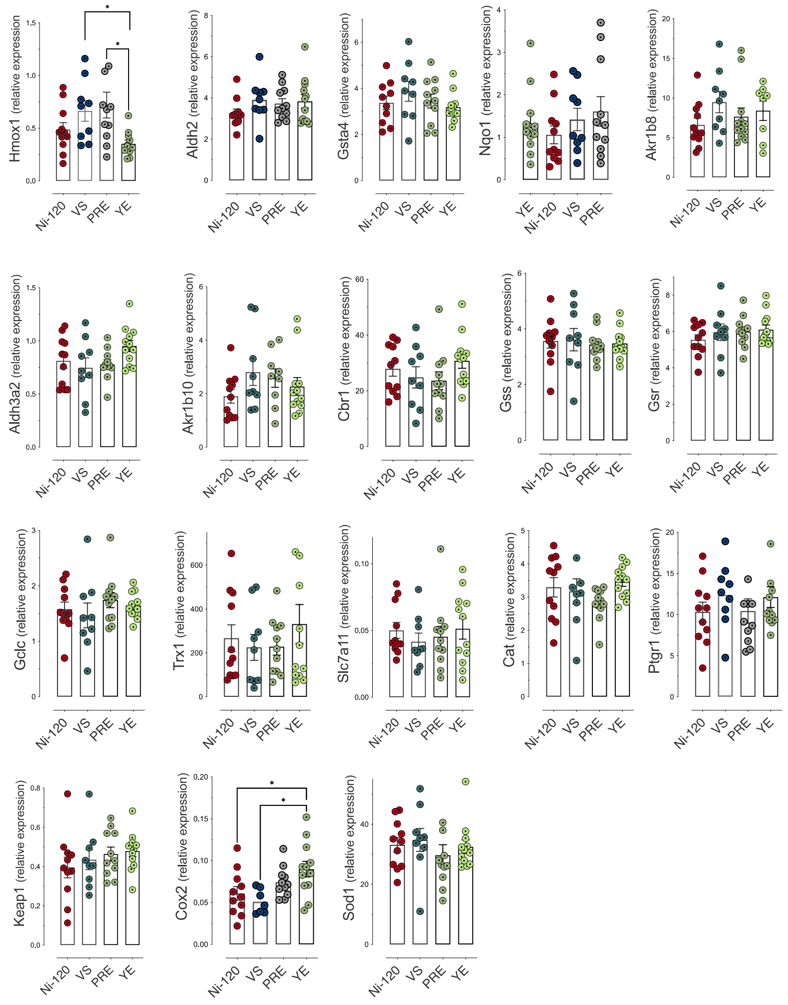


**Figure S5. sodium nitrite alternative effect (Vegetable Stock (VS), Polyphenol-rich Extract (PRE), Lallemand solution (YE) on gene expression in rat colon mucosa.** Data were represented using scatter plots with bar (mean ± sem), * $p \leq 0.05$. Genes full names are shown in Supplementary Table S9.

### 6. *Assessment of 8-isoprostane (8-isoPGF$_{2\alpha}$) in urinary samples of the 100 days-study with different level of sodium nitrite (0, 90 and 120 mg/kg) or alternatives to sodium nitrite in cooked ham models.*

#### *6.1 Supplementary Experimental Section*

8-isoPGF$_{2\alpha}$ was assayed using Cayman ELISA kits (Bertin Technologies), according to the manufacturer's instructions.

#### *6.2 Supplementary Results*

Modification of the sodium nitrite levels (Figure S6A) or sodium nitrite substitution by alternatives (Figure S6B) had no effect of urinary excretion of 8-isoPGF$_{2\alpha}$ per 24h.

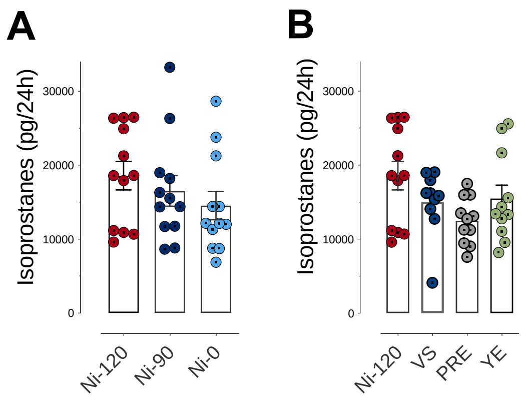


**Figure S6. Impact of sodium nitrite concentrations and alternatives on urinary 8-isoPGF$_{2\alpha}$ (pg/24h).**

**A-**Impact of sodium nitrite concentrations in processed meats (0 *vs* 90 *vs* 120 mg/kg). **B-** Impact of alternatives (Vegetable Stock (VS), Polyphenol-rich Extract (PRE), Lallemand solution (YE)). Data were represented using scatter plots with bar (mean ± sem).

#### 7. *Abundance variation of fecal bacterial communities displaying dose effects and/or changes in response to alternatives to sodium nitrite in cooked ham model.*

##### *7.1* Table S10

**A-**Taxonomic affiliation of clusters agglomerated at the genus rank significantly affected by the sodium nitrite content in ham-based diets (Deseq2, Padj≤0.05).

| OTU | baseMean | log2FoldChange | lfcSE | stat | pvalue | padj | Phylum | Class | Order | Family | Genus |
|---|---|---|---|---|---|---|---|---|---|---|---|
| Cluster_16 | 1207.38 | 1.74533 | 0.426143 | 20.8113 | 0.0000303 | 0.00164212 | Firmicutes | Clostridia | Lachnospirales | Lachnospiraceae | unknown genus |
| Cluster_144 | 26.4446 | 1.2813 | 0.52824 | 20.2749 | 0.0000396 | 0.00164212 | Firmicutes | Clostridia | Peptococcales | Peptococcaceae | unknown genus |
| Cluster_929 | 2.92678 | 2.71891 | 0.901342 | 17.9768 | 0.000124851 | 0.0034542 | Firmicutes | Clostridia | Lachnospirales | Lachnospiraceae | Eisenbergiella |
| Cluster_5 | 2449.29 | -0.930625 | 0.500487 | 15.1397 | 0.000515762 | 0.0107021 | Firmicutes | Clostridia | Peptostreptococcales-Tissierellales | Peptostreptococcaceae | Romboutsia |
| Cluster_122 | 24.2941 | -1.40029 | 0.435521 | 13.0645 | 0.00145574 | 0.0241653 | Firmicutes | Clostridia | Oscillospirales | Ruminococcaceae | Candidatus Soleaferrea |
| Cluster_96 | 46.8693 | -0.0712521 | 0.542587 | 9.77815 | 0.00752839 | 0.0414277 | Firmicutes | Clostridia | Lachnospirales | Lachnospiraceae | [Ruminococcus] torques group |

**B-**Taxonomic affiliation of clusters agglomerated at the genus rank significantly affected by the alternative to nitrites in ham-based diets (Deseq2, Padj≤0.05).

| OTU | baseMean | log2FoldChange | lfcSE | stat | pvalue | padj | Phylum | Class | Order | Family | Genus |
|---|---|---|---|---|---|---|---|---|---|---|---|
| Cluster_31 | 366,638 | -0,927404 | 0,49264 | 124,185 | 9,68E-27 | 8,04E-25 | Firmicutes | Clostridia | Lachnospirales | Lachnospiraceae | GCA-900066575 |
| Cluster_161 | 18,2941 | -3,78975 | 0,655325 | 40,4462 | 8,57E-09 | 3,56E-07 | Firmicutes | Bacilli | Erysipelotrichales | Erysipelotrichaceae | Faecalitalea |
| Cluster_113 | 38,5171 | 2,48452 | 0,51532 | 38,3481 | 2,39E-08 | 6,60E-07 | Bacteroidota | Bacteroidia | Bacteroidales | Muribaculaceae | Muribaculum |
| Cluster_16 | 1 556,47 | 2,46239 | 0,447278 | 35,2795 | 1,06E-07 | 0,00000177 | Firmicutes | Clostridia | Lachnospirales | Lachnospiraceae | unknown genus |
| Cluster_133 | 11,1148 | -6,60138 | 1,24078 | 35,4868 | 9,61E-08 | 0,00000177 | Firmicutes | Clostridia | Oscillospirales | Oscillospiraceae | Flavonifractor |
| Cluster_416 | 6,01094 | -0,799438 | 0,618033 | 34,7018 | 1,41E-07 | 0,00000195 | Bacteroidota | Bacteroidia | Bacteroidales | Prevotellaceae | Prevotellaceae UCG-001 |
| Cluster_64 | 41,625 | 1,23016 | 0,80329 | 26,9295 | 0,00000609 | 0,0000722 | Actinobacteriota | Actinobacteria | Bifidobacteriales | Bifidobacteriaceae | Bifidobacterium |
| Cluster_93 | 46,0782 | -1,62564 | 0,545465 | 21,3623 | 0,0000885 | 0,000918454 | Firmicutes | Clostridia | Oscillospirales | Ruminococcaceae | Anaerotruncus |
| Cluster_6 | 1 698,23 | -0,612995 | 0,184828 | 18,4109 | 0,000361835 | 0,00333692 | Desulfobacterota | Desulfovibrionia | Desulfovibrionales | Desulfovibrionaceae | Bilophila |
| Cluster_15 | 689,674 | 0,201521 | 0,229197 | 14,9406 | 0,0018681 | 0,0140956 | Proteobacteria | Gammaproteobacteria | Burkholderiales | Sutterellaceae | Parasutterella |
| Cluster_3 | 2 124,80 | 1,49192 | 0,939298 | 15,0091 | 0,00180887 | 0,0140956 | Firmicutes | Clostridia | Clostridiales | Clostridiaceae | Clostridium sensu stricto 1 |
| Cluster_5 | 1 659,01 | -0,931654 | 0,612477 | 14,0093 | 0,00289248 | 0,0200063 | Firmicutes | Clostridia | Peptostreptococcales-Tissierellales | Peptostreptococcaceae | Romboutsia |
| Cluster_96 | 38,3441 | -1,38387 | 0,512519 | 13,2188 | 0,00418655 | 0,0267295 | Firmicutes | Clostridia | Lachnospirales | Lachnospiraceae | [Ruminococcus] torques group |
| Cluster_122 | 36,7519 | -1,05584 | 0,434465 | 12,3789 | 0,00619171 | 0,036708 | Firmicutes | Clostridia | Oscillospirales | Ruminococcaceae | Candidatus Soleaferrea |
| Cluster_130 | 19,3984 | 1,03016 | 0,553244 | 10,9901 | 0,0117793 | 0,0551788 | Firmicutes | Clostridia | Lachnospirales | Lachnospiraceae | Tuzzerella |

#### *7.2 Supplementary Figure*

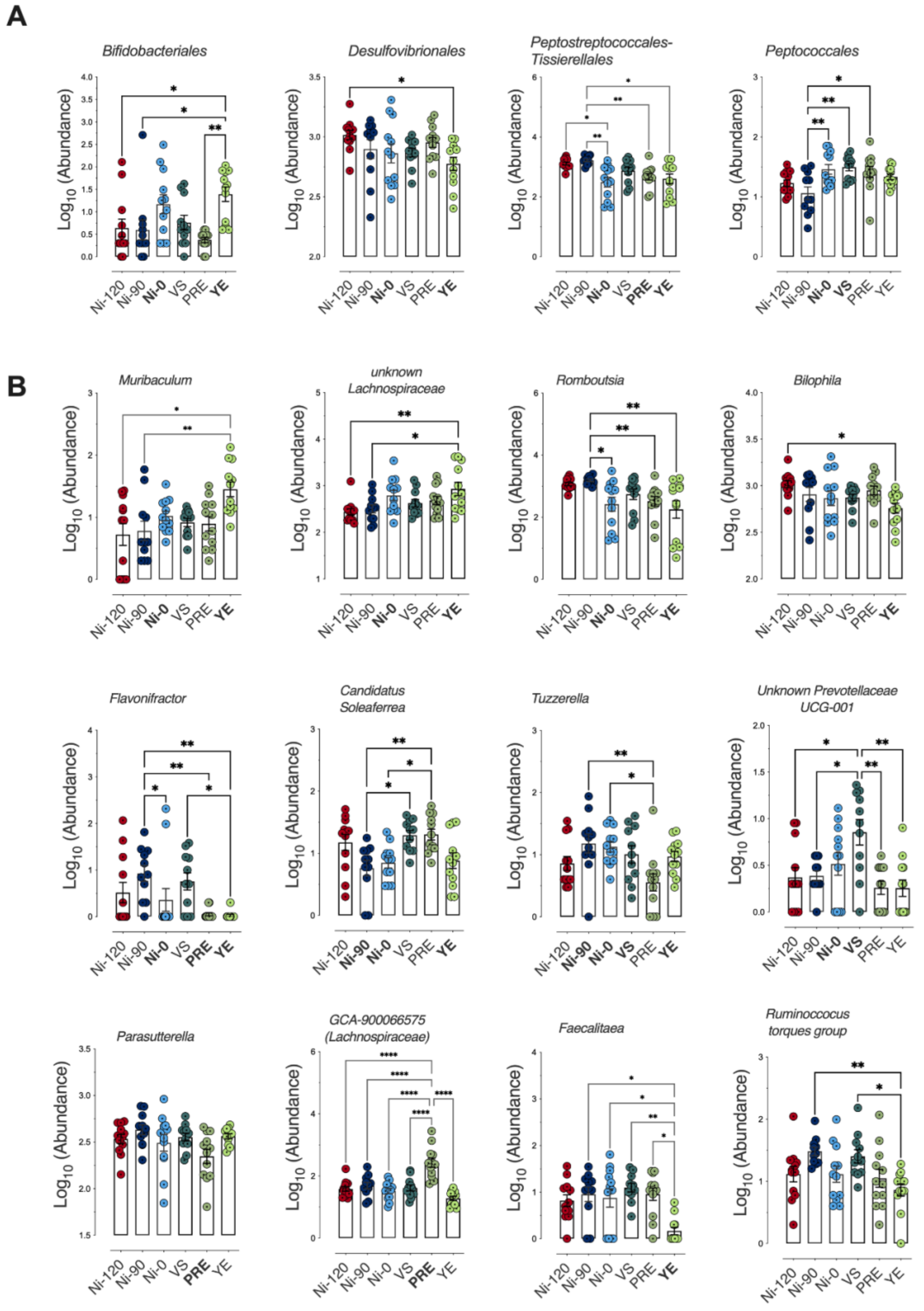


**Figure S7. Impact of sodium nitrite concentrations and sodium nitrite alternatives on fecal microbiota of rats.** Normalized $Log_{10}$ abundances at the order level (**A**) and genus level (**B**) resulted from agglomeration of OTUs. Data were represented using scatter plots with bar (mean ± sem). * $p \leq 0.05$; ** $p \leq 0.01$, ** $p \leq 0.01$, *** $p \leq 0.001$, **** $p \leq 0.0001$.
